# A comprehensive proteomic SWATH-MS workflow for profiling blood extracellular vesicles: a new avenue for glioma tumour surveillance

**DOI:** 10.1101/2020.03.05.979716

**Authors:** Susannah Hallal, Ali Azimi, Heng Wei, Nicholas Ho, Maggie Lee, Hao-Wen Sim, Joanne Sy, Brindha Shivalingam, Michael E. Buckland, Kimberley L. Kaufman

**Affiliations:** Neurosurgery Department, Chris O’Brien Lifehouse, Camperdown, NSW; Brainstorm Brain Cancer Research, Brain and Mind Centre, The University of Sydney, NSW; Discipline of Pathology, School of Medical Sciences, The University of Sydney, NSW; Dermatology Department, School of Medicine and Health, The University of Sydney, Westmead NSW; Neuropathology Department, Royal Prince Alfred Hospital, Camperdown NSW; Department of Medical Oncology, Chris O’Brien Lifehouse, Camperdown, NSW; NHMRC Cinical Trials Centre, University of Sydney, Camperdown, NSW; The Kinghorn Cancer Centre, St Vincent’s Hospital, Darlinghurst, NSW

**Keywords:** Glioblastoma, Glioma, Extracellular Vesicle, Plasma, SWATH, DIA, Data independent acquisition, mass spectrometry

## Abstract

There is a real need for biomarkers that can indicate glioma disease burden and inform clinical management, particularly in the recurrent glioblastoma (GBM; grade IV glioma) setting where treatment-associated brain changes can confound current and expensive tumour surveillance methods. In this regard, extracellular vesicles (EVs; 30-1000 nm membranous particles) hold major promise as robust tumour biomarkers. GBM-EVs encapsulate molecules that reflect the identity and molecular state of their cell-of-origin and cross the blood-brain-barrier into the periphery where they are readily accessible. Despite the suitability of circulating-EVs for GBM biomarker discovery, sample complexity has hindered comprehensive quantitative proteomic studies. Here, sequential window acquisition of all theoretical fragment ion spectra mass spectrometry (SWATH-MS) was used in conjunction with a targeted data extraction strategy to comprehensively profile circulating-EVs isolated from plasma. Plasma-EVs sourced from pre-operative glioma II-IV patients (*n*=41) and controls (*n*=11) were sequenced by SWATH-MS, and the identities and absolute quantities of the proteins were extracted by aligning the SWATH-MS data against a custom glioma spectral library comprised of 8662 high confidence protein species. Overall, 4054 plasma-EV proteins were quantified across the cohorts, and putative circulating-EV biomarker proteins identified (*adjusted* p-value<0.05) included previously reported GBM-EV proteins identified *in vitro* and in neurosurgical aspirates. Principle component analyses showed that plasma-EV protein profiles clustered according to glioma subtype and WHO-grade, and plasma-EV proteins reflected the extent of glioma aggression. Using SWATH-MS, we describe the most comprehensive proteomic plasma-EV profiles for glioma and highlight the promise of this approach as an accurate and sensitive tumour monitoring method. Objective blood-based measurements of glioma tumour activity will support the implementation of next-generation, patient-centred therapies and are ideal surrogate endpoints for recurrent progression that would allow clinical trial protocols to be more dynamic and adapt to the individual patient and their cancer.

## Introduction

Diffuse gliomas are the most common and devastating primary malignant brain tumours of adults, owing to their characteristic diffusely invasive growth patterns. The principal histologic subtypes, astrocytoma and oligodendrogliomas, are graded by severity according to a set of prescribed histological and molecular features, ranging from WHO grade II-IV (least to most severe), with both subtype and grade contributing to prognosis [1]. The most prominent distinguishing molecular feature of diffuse gliomas is the isocitrate dehydrogenase (*IDH*) mutational status, which separates tumours into two broad categories. In general, *IDH*-wildtype astrocytic tumours arise as primary *de novo* glioblastomas (GBM; astrocytoma grade IV) and are the most frequent and aggressive manifestations of diffuse glioma. GBMs are almost universally fatal and carry a dismal median survival of 15 months [2]. The current standard of care for GBM patients involves debulking surgery to remove the main tumour mass, followed by concomitant radiotherapy and temozolomide (TMZ) chemotherapy (STUPP protocol) [3]. Unfortunately, treatments only provide temporary palliation to patients, and GBM recurrences are inevitable [4]. GBM tumours often re-emerge as evolved, treatment-resistant entities and death typically ensues within 6-months of recurrence [2]. *IDH* mutations confer a prognostic advantage and are diagnostic of grade II-III gliomas or secondary GBMs that have progressed from grade II-III astrocytomas [5, 6]. *IDH*-mut tumours are further subdivided according to 1p/19q codeletion status, where oligodendrogliomas harbor both *IDH*-mut and 1p/19q codeletion and carry the most favourable prognosis. Malignant progression of *IDH*-mutated gliomas is also common, including the appearance of a ‘hypermutation phenotype’ following TMZ therapy [7].

Apart from a lack of targeted treatment strategies, a significant obstacle to effective clinical management of diffuse glioma is the dearth of sensitive approaches to monitor tumour progression and/or treatment-resistant recurrence. Histological evaluation of brain tissue is the only definitive method for diagnosing glioma progression and/or recurrence [8]. Yet routine neurosurgical biopsies are impractical for tumour surveillance, contribute significantly to patient morbidity [9] and have inherent under-sampling issues due to the highly heterogenous nature of these tumours [10]. Neuroimaging and neurological clinical assessments are the current mainstay for monitoring glioma tumours, but often pose challenges to accurate tumour monitoring. Despite significant technological advances, neuroimaging approaches are insensitive to early signs of recurrence. Tumour progression can also be confounded by treatment-related effects, such as pseudoprogression and radiation necrosis, as they share radiological features with tumour recurrence [11-17]. Generally, pseudoprogression is observed as a radiographic worsening of the tumour and is indistinguishable from true tumour progression [15]. A meta-analysis of 73 studies reviewed MRI progression in 2603 high-grade glioma patients and determined a 36% incidence rate of pseudoprogression [13]. Despite new and promising advanced imaging techniques, pseudoprogression remains indistinguishable from early tumour progression [15]. Radionecrosis is a severe local tissue inflammatory reaction in response to radiotherapy [16] and shares considerable radiographic features with tumour recurrence [17]; there is often a combination of both entities [18] and such cases are notoriously difficult to diagnose and manage [17]. While revisions have been made to Response Assessment in Neuro-Oncology (RANO) criteria to standardise radiographic tumour monitoring [19], it remains challenging to accurately assess treatment-related effects from true tumour growth. In some cases, therapy is administered with the uncertainty of whether aggressive treatment is warranted, while other patients with true tumour progression are treated with a ‘wait and see’ approach rather than being switched to potentially life-extending treatment. Although understudied, this extended period without a definitive diagnosis adds to the already heavy physiological and psychological burden borne by patients and their care givers [20].

Improving patient outcomes ultimately requires the development of sensitive methods that can accurately, efficiently and sensitively monitor glioma activity and treatment response. With efforts to improve the clinical management of cancer comes a growing trend to design minimally-invasive liquid biopsies that can better diagnose, monitor and guide treatment decisions [21-23]. Liquid biopsies measure tumour-derived factors in body fluids, offering ease of accessibility to tumour-molecules and the added advantage of monitoring tumour evolution in real-time [21]. Such ready access to glioma molecular information would provide clinicians much-needed information to better guide treatment, prevent unnecessary neurosurgeries and allow treatments to be better synchronised for maximum therapeutic benefit [21]. Currently, there are no such glioma blood- or CSF-based biomarkers. The development of a glioma liquid biopsy would require specific biomarkers that are stable and easily accessible from body fluids and reflect the identity and molecular state of the tumour, to which extracellular vesicles (EVs) hold major promise. EVs are 30-1000 nm membranous particles that are released by all cell types. We have shown that glioma-EVs can be sampled directly from the glioma microenvironment and are capable of stratifying patients [24, 25]. EV secretion is increased in neoplasia [26, 27] and glioma-derived EVs cross the blood-brain-barrier (BBB) into the circulation [28, 29], allowing peripheral sampling of glioma molecular information [30]. Capturing glioma-EVs from body fluids and screening their molecular contents may therefore serve as a complementary approach to assess the heterogeneic molecular landscape of gliomas in real-time as tumours evolve [21].

Despite the suitability of circulating-EVs for biomarker discovery, comprehensive proteomic characterisation has been challenging. Glioma-EVs constitute only a minor subset of the total circulating-EV population, with the majority being derived from platelets and endothelial cells [31]. While attempts have been made to capture and analyse pure circulating GBM-EVs by targeting common GBM cell-membrane proteins [32], no universal or ‘pan’ GBM or glioma-EV surface markers have been identified that can accurately and reproducibly target circulating GBM-EVs. Furthermore, cell-membrane proteins are topologically reversed on the EV surface [33]; thus the unequivocal and reliable capture of circulating GBM-EVs for targeted biomarker investigations requires comprehensive characterisations of GBM-EV surface proteomes. Yet, total circulating-EV populations may reflect the systemic effects of tumour pathology and/or radio-chemotherapy [34] and analysing only a subset of circulating-EVs might omit systemic effects that could otherwise indicate disease burden and treatment effectiveness.

Perhaps the most significant factor that has hindered large-scale proteomics of circulating-EVs is the complexity of the blood proteome. Despite recent advancement in mass spectrometry (MS) and blood-EV isolation methods, the sheer complexity of the blood, with an array of protein species of varying abundance, remains a major obstacle to large-scale proteomic analyses. Albumin, a large transport protein, is reported to be the most abundant protein within the blood (55%) and a wide dynamic range in abundance is reported for all other proteins [35]. Albumin forms a complex network with multiple proteins, lipids and peptide hormones, known as the ‘albuminome’ [36, 37], making its selective, specific depletion challenging. In addition to albumin, other highly-abundant blood proteins such as haemoglobin, serotransferrin, complement and immunoglobulins commonly isolate with blood-EVs [38]. These highly-abundant proteins mask the detection of less-abundant, potential EV biomarker proteins in traditional *shot-gun* MS analyses [39] based on information dependent acquisition (IDA) methods [40]. The limitation of IDA in biomarker discovery is due to its inherent dependence on inclusion lists which only allow highly abundant peptides to be assessed by MS. The complexity of the blood has served as a major limitation to IDA-based biomarker discovery analyses, as putative disease-related biomarkers often go undetected, unidentified and neglected [39].

Due to limitations in the current discovery-based proteomic approaches, more recent MS strategies rely on highly-specific data independent acquisition (DIA) methods for targeted analysis of proteins within complex biological mixtures. Sequential window acquisition of all theoretical mass spectra (SWATH) is a form of DIA on Sciex TripleTOF 5600+ instruments. SWATH is a label-free MS method that theoretically allows all peptides in a sample to be identified and quantified. The method was first described by Gillet *et al* [41] and involves a targeted data extraction strategy that mines SWATH-MS data against an IDA spectral library [42]. The SWATH-MS method involves fragmenting and analysing all ionised peptides across SWATH windows of a specified mass-range in an unbiased fashion so that all ions undergo MS/MS, enabling sensitive and accurate quantitation, even for low abundant peptides [43, 44]. High-resolution extracted ion chromatograms (XICs) are drawn for the fragment ions for every peptide in the sample [45]. The SWATH-MS data can then be aligned to a high-quality comprehensive spectral library containing MS coordinates for each target peptide: (i) the peptide precursor ion *m/z*, (ii) the *m/z* of the fragment ions and their intensities and (iii) the chromatographic retention time of the peptide [45]. This information allows proteins that were detected by SWATH-MS to be identified and quantified if they are present in the library [45]. As such, SWATH-MS data can be archived and analysed retrospectively, allowing for maximal identifications as spectral libraries mature.

This study aimed to profile the proteomic contents of circulating-EVs from glioma patients to determine a set of putative markers that are indicative of glioma subtype and progression. Here, a targeted SWATH-MS proteomic workflow was utilised for the absolute identification and quantification of plasma-EV proteins. A comprehensive spectral library comprised of peptides derived from EV, GBM and other tumour specimens was used to extract proteins detected in circulating-EVs by SWATH-MS. As discoveries in circulating-EVs have immense potential for the development of new clinical tests, it was important to optimise the experimental workflow with an EV isolation method, that isolates relatively pure EV populations in a rapid, efficient and scalable fashion, i.e., size exclusion chromatography (SEC), so that future EV biomarker panel tests can be readily adopted by hospital clinical pathology services. We show that SWATH-MS is a suitable and promising method for advancing biomarker discovery using EVs derived from complex biological fluids, such as the blood. Plasma-EV proteomes were able to stratify glioma patients and EVs sampled from recurrent and progressed tumours showed protein changes consistent with more aggressive glioma-EV profiles. These exciting findings may be pivotal for the shift towards precision care models in the management of diffuse glioma.

## Materials and Methods

### Pre-operative plasma cohorts

Pre-operative bloods were collected from patients prospectively over a 36-month period. All patients provided research consent (Royal Prince Alfred Hospital Neuropathology Tumour and Tissue Bank, SLHD HREC protocol X14-0126) and specimens were processed and analysed under approved University of Sydney HREC project 2012/1684. Diagnoses were made by neuropathological assessment of surgically-resected tumour material. Bloods were sampled from patients with primary GBM (grade IV *IDH-*wt astrocytoma, *n*=24), secondary GBM (grade IV *IDH-*mut astrocytoma, *n*=2), glioma grade II-III [*IDH-*mut astrocytoma (AST), *n*=12; *IDH-*mut, 1p19q codeleted oligodendroglioma (OLI; *n*=4)] and meningioma grade I (MEN, *n*=5; non-glioma control). Within this patient cohort, bloods were re-sampled (prior to repeat surgeries) from three individual patients with confirmed glioma recurrence. Matching, re-sampled blood was captured from a patient with: 1) progression from grade II to grade III *IDH-*mut astrocytoma; 2) progression from grade III to grade IV *IDH-*mut astrocytoma; and 3) recurrence of primary *IDH-*wt GBM. Plasma from healthy age-gender matched controls (HC, *n*=6; 3 males, 3 females, age range 38-55 years) were also included. A summary of sample cohorts used for comparative analyses is provided in Table 1. A more detailed summary of patient demographics and diagnoses are provided in Supplementary Table 1.

**Table 1.** Overview of pre-operative blood specimen cohorts and grouping schema. Plasma-EV samples were grouped by their respective genetic-histological subtype and WHO grade.

| **Genetic-histological subtype** |  |  |  |  |
| --- | --- | --- | --- | --- |
| Cohort | Sample (*n*) | Gender | Age at resection (Mean ± SD) | Histopathological diagnosis |
| GBM | 24 | 17M/7F | 60.8 ± 14.1 | Primary Glioblastoma, *IDH*-wildtype, WHO grade IV |
| AST | 13 | 10M/3F | 40.1 ± 13.8 | Astrocytoma, *IDH*-mutant; WHO grade II (*n* = 5), grade III (*n* = 6) and grade IV (*n* = 2) |
| OLI | 4 | 4M | 36.5 ± 3.7 | Oligodendroglioma, *IDH*-mutant; WHO grade II (*n* = 1) and grade III (*n* = 3) |
| **WHO Grade** |  |  |  |  |
| Cohort | Sample (*n*) | Gender | Age at resection (Mean ± SD) | Histopathological diagnosis |
| GIV | 26 | 19M/7F | 59.4 ± 14.6 | WHO grade IV astrocytoma; *IDH*-wildtype [*n*=24] and *IDH*-mutant Glioblastoma [*n*=2] |
| GIII | 9 | 8M/1F | 39.8 ± 13.5 | WHO grade III glioma; *IDH*-mutant astrocytoma and *IDH*-mutant oligodendroglioma |
| GII | 6 | 4M/2F | 37.3 ± 11.9 | Grade II glioma; *IDH*-mutant astrocytoma and *IDH*-mutant oligodendroglioma |
| **Controls** |  |  |  |  |
| Cohort | Sample (*n*) | Gender | Age at resection (Mean ± SD) | Histopathological diagnosis |
| MEN | 5 | 3M/2F | 57.6 ± 11.0 | Meningioma, grade I<br>**Non-glioma control** |
| HC | 6 | 3M/3F | 46.8 ± 7.4 | Healthy controls |

### Plasma processing and EV isolation

Peripheral blood (15 mL) was collected into BD Vacutainer K2 EDTA tubes containing spray-dried K2 EDTA and processed within 4 h. The plasma was separated by a Ficoll-histopaque gradient (400 x *g*, 30 min, RT; no brake). Platelets were depleted from the plasma by centrifugation (3000 x *g*, 20 min, 4 °C) and 1.5 mL plasma aliquots were snap frozen in liquid N_2_ and stored at −80 °C.

Platelet-free plasma was thawed slowly on ice and EVs were purified using iZON qEV_original_ columns as per manufacturer’s instructions. Briefly, qEV columns were equilibrated with 5 mL PBS before loading 500 µL of platelet-free plasma onto the column. When the plasma passed through the reservoir, twelve 500 µL fractions were eluted in PBS. While each column can perform up to five EV isolations with PBS washes in between samples, only one column was used per specimen to circumvent signal carryover or cross contamination. The plasma-EV fractions 1-12 were stored at −80°C.

### Preparation of EV proteomes for LC-MS/MS

The volume of plasma-EV fractions was reduced by vacuum centrifugation to 100 µL and EV proteins precipitated using chloroform-methanol. The precipitated EV protiens were dissolved in 90% (v/v) formic acid (FA) and dried by vacuum centrifugation. The dried EV proteins were resuspended in 0.2% (wt/vol) Rapigest SF™ (Waters, Milford, MA) in 0.05 mol/L triethylammonium bicarbonate (TEAB) and incubated at 95 °C for 5 min. Samples were sonicated twice with a step-tip probe at 30% intensity for 30 seconds to aid protein resuspension. Debris was removed by centrifugation at 13,000 rpm for 20 min at 4 °C. Proteins were then reduced in 12 mM Tris (2-carboxyethyl) phosphine (TCEP) (60 °C, 30 min) and alkylated in 50 mM iodoacetamide (RT, 30 min, in the dark). Sample pH was adjusted to 7.5 with 0.05 M TEAB and proteins were digested overnight at 37 °C with sequencing-grade trypsin (Promega, Madison, WI) in 1:30 (w/w) trypsin:protein ratio. The peptide solution was adjusted to a pH<2 using 50% (v/v) FA and incubated for 30 min at 37 °C to cleave the Rapigest SF™ detergent, which was then removed by centrifugation (16,000 x *g*, 10 min, 4 °C). Peptides were then desalted by solid-phase extraction using 1cc HLB cartridges (Waters, MA, USA) and eluted in 70% acetonitrile (ACN)/0.1% FA (v/v). Protein and peptide concentrations were measured in duplicate using the Qubit^®^ Protein Assay Kit (Invitrogen, Carlsbad, CA). Peptides were dried by vacuum centrifugation in 25 μg aliquots. The dried, ‘clean’ peptide mixtures were stored at −80 °C.

### EV characterisation by Nanoparticle Tracking Analysis, Transmission Electron Microscopy and LC-MS/MS

The size distributions and concentrations of the plasma-EV fractions were measured by nanoparticle tracking analysis (NTA) software (version 3.0) in triplicate using the NanoSight LM10-HS (NanoSight Ltd, Amesbury, UK), configured with a tuned 532 nm laser and a digital camera system (CMOS trigger camera). EVs were diluted with sterile-filtered PBS (viscosity 1.09 cP) to ensure 20-100 particles were detectable within the field of view of the standard CCD camera of the microscope. The NTA software captured triplicate 60 s video recordings of the EVs, at 25 frames per second with default minimal expected particle size, minimum track length, blur setting, with the temperature of the laser unit controlled to 25 °C. The videos were analysed by NTA3.0, which translates the Brownian motion and light scatter properties of each individual laser-illuminated particle into a size distribution (ranging from 10 to 1000 nm) and concentration (particles per mm) while simultaneously calculating their diameter using statistical methods [46]. The EV size distributions and concentrations were analysed in Microsoft Excel^®^. Plasma-EVs were also analysed by transmission electron microscopy (TEM). EVs were re-suspended in dH_2_O, loaded onto carbon-coated, 200 mesh Cu formvar grids (ProSciTech Pty Ltd; Cat No. GSCU200C) and fixed with 2.5% glutaraldehyde in 0.1 M phosphate buffer (pH 7.4). Samples were negatively stained with 2% uranyl acetate for 2 min, dried at RT for 3 h and then visualised at 40,000 X, 80,000 X and 100,000 X magnification on a Philips CM10 Biofilter TEM (FEI Company, OR, USA) equipped with an AMT camera system (Advanced Microscopy Techniques, Corp., MA, USA) at an acceleration voltage of 80-120 kV.

The presence of canonical EV-marker proteins in plasma-EV fractions was confirmed by liquid chromatography tandem mass spectrometry (LC-MS/MS) using a Q-Exactive plus hybrid quadrupole-orbitrap mass spectrometer (Thermo Scientific, MA, USA). Peptide mixtures resuspended to 66.7 ng/µL in 3% (v/v) ACN/0.1% (v/v) FA were separated by nano-LC using an Ultimate 3000 UHPLC and autosampler system (Dionex, Amsterdam, Netherlands). Reverse-phase mobile buffers were composed of A: 0.1% (v/v) FA (Thermo Scientific, MA, US, Cat No. 34851-4), and B: 80% (v/v) ACN (Thermo OPTIMA LC-MS grade, Cat No. 34851-4) /0.1% FA. Peptides were eluted using a linear gradient of 5% B to 42% B over 120 min with a constant flow rate of 250 nL min^-1^. High voltage (2000 V) was applied to a low volume tee (Valco, Houston, TX, USA) and the column tip positioned approximately 0.5 cm from the heated capillary (T = 275 °C) of the mass spectrometer. Positive ions were generated by electrospray and the Orbitrap was operated in IDA mode. A survey scan of 350-1550 *m/z* was acquired. Up to ten of the most abundant ions (>5000 counts) with charge states ≥ +2 were selected and fragmented by collision induced dissociation (CID; activation time of 10 ms). Mass-to-charge (*m/z*) ratios selected for MS/MS were dynamically excluded for 20 s. Prior to loading the samples, an LC-MS/MS standard consisting of 30 fmol pre-digested bovine serum albumin (BSA; GeneSearch, QLD, Australia, Cat No. P8108S, 500 pmol) was injected to ensure optimal performance and dynamic range of the instrument. Raw MS/MS results were analysed by Thermo Proteome Discoverer (version 2.2.0.388), followed by a MS/MS ion search of peak lists for all MS/MS samples using Mascot (Matrix Science, London, UK; version 2.4.0) against the SwissProt database with the following parameters: i) trypsin, up to two missed cleavages; ii) taxonomy, *Homo sapiens*; iii) no fixed peptide modifications; (iv) variable peptide modifications, carbamidomethylation (C), and oxidation (M); (v) peptide tolerance ± 4 ppm; (vi) MS/MS tolerance ± 0.1 Da; (vii) Peptides with charge 2+, 3+, and 4+; and (viii) instrument ESI–FT ICR (fourier-transform ion cyclotron resonance). Mascot .dat files were imported into Scaffold (version 4.8.7, Proteome Software Inc, OR, USA) and analysed against the SwissProt Database (SwissProt_2018_05) with X! Tandem (The GPM, thegpm.org; version CYCLONE (2010.12.01.1). Mascot and X! Tandem were searched with a fragment ion mass tolerance of 0.10 Da and a parent ion tolerance of 4.0 ppm. Oxidation of methionine and carbamidomethylation of cysteine were specified in X! Tandem as variable modifications. Peptide and protein identifications were validated in Scaffold 4 (Proteome Software, Portland, Oregon) with the peptide [47] and protein [48] prophet algorithms, respectively. Peptide identifications were accepted if they could be established at greater than 95.0% probability with scaffold delta-mass correction, while protein identifications were accepted if they could be established at greater than 99.0% probability with at least two peptides identified at 95.0% or greater. False discovery rate (FDR) thresholds were calculated by searching the data against a decoy database. Proteins were grouped with protein cluster analysis. Proteins that contained similar peptides and could not be differentiated based on MS/MS analysis alone were grouped to satisfy the principles of parsimony [49]. Protein identity ambiguity was removed by manually deselecting all peptides shared by more than one protein. Protein abundances were exported to Microsoft Excel^®^ as normalised total spectra for analysis.

### Hydrophilic Interaction Liquid Chromatography (HILIC)

A range of GBM samples (EVs, cells and tumour tissue fractions) were used to generate a comprehensive, custom glioma IDA spectral library. Specimens were prepared as described before from neurosurgical aspirate EVs [24], patient-derived GBM-stem-like cells [50], and soluble and microsomal proteomes [51] prepared from GBM tumour tissue (Supplementary Table 2). The peptide samples were fractionated by HILIC, a reverse-phase chromatographic method, to reduce sample complexity and increase protein ID coverage by LC-MS/MS [52]. Briefly, desalted pooled peptides (20 µg) were dried by vacuum centrifugation, resuspended to 0.5 µg/µL in 90% ACN/0.1% trifluoroacetic acid (TFA) and fractionated with an Agilent 1200 high performance liquid chromatography (HPLC) system (Agilent Technologies, Santa Clara, CA) on an in-house 17 cm TSKgel Amide-80 HILIC column (3 µm particle size) with a post-column *PEEK*™ filter (Upchurch Scientific, Rohnert Park, CA, USA). Mobile phase buffers were comprised of A: 0.1% (v/v) TFA (Sigma, Chromasolv HPLC grade, Cat.No. 34851-4) and B: 90% (v/v) ACN/0.1% (v/v) TFA. The samples were loaded in buffer B and eluted over a 50 min gradient comprised of 100% B for 10.5 min (flow rate of 10 µL/min), followed by 100-60% B in 26.5 min, 60-30% B in 4 min, 30% B for 1 min and 100% A for the rest of the gradient at a flow rate of 6 µL/min. Chromatographic performance was monitored at 210 nm. Uniform peptide fractions were eluted in buffer B between 13-35 min. The number of HILIC fractions collected is specified in Supplementary Table 2. The resultant peptide fractions were dried by vacuum centrifugation.

### IDA and SWATH-MS Analysis

LC-MS/MS data acquisition (IDA and SWATH) was performed on an AB Sciex TripleTOF^®^ 6600 Quadrupole Time-Of-Flight (QTOF) mass analyser. The TripleTOF^®^ system was operated in IDA mode to acquire the spectral library data for HILIC fractionated peptides (Supplementary Table 2). SWATH-MS mode was utilised to analyse the plasma-EV proteomes (Table 1). The workflow is presented in Figure 1. Dried peptides were resuspended in 3% ACN (v/v)/ 0.1% (v/v) FA to a concentration of 0.4 µg/µL and *PepCalMix* heavy-labelled peptides (AB Sciex) were added to each fraction to a final concentration of 6.67 fmol/µL. The resuspended peptides (2 µg) were injected onto an in-house 15 cm C_18_ reversed-phase column (75 µm diameter and 5 µm particle size) and analysed by the TripleTOF^®^6600 coupled to an Eskpert™ NanoLC 425. The HPLC solvent system was comprised of buffer A: 0.1% (v/v) FA (Thermo Scientific, Cat.No. 85178) and buffer B: 80% (v/v) ACN (Thermo OPTIMA LC/MS grade, Cat.No. 34851-4), 0.1% (v/v) FA. Peptides were eluted over a 120 min gradient (2%-35% B for 80 min, 35-95% B for 19 min, 95% B for 5 min and 2% B for 16 min).

**Figure 1.**
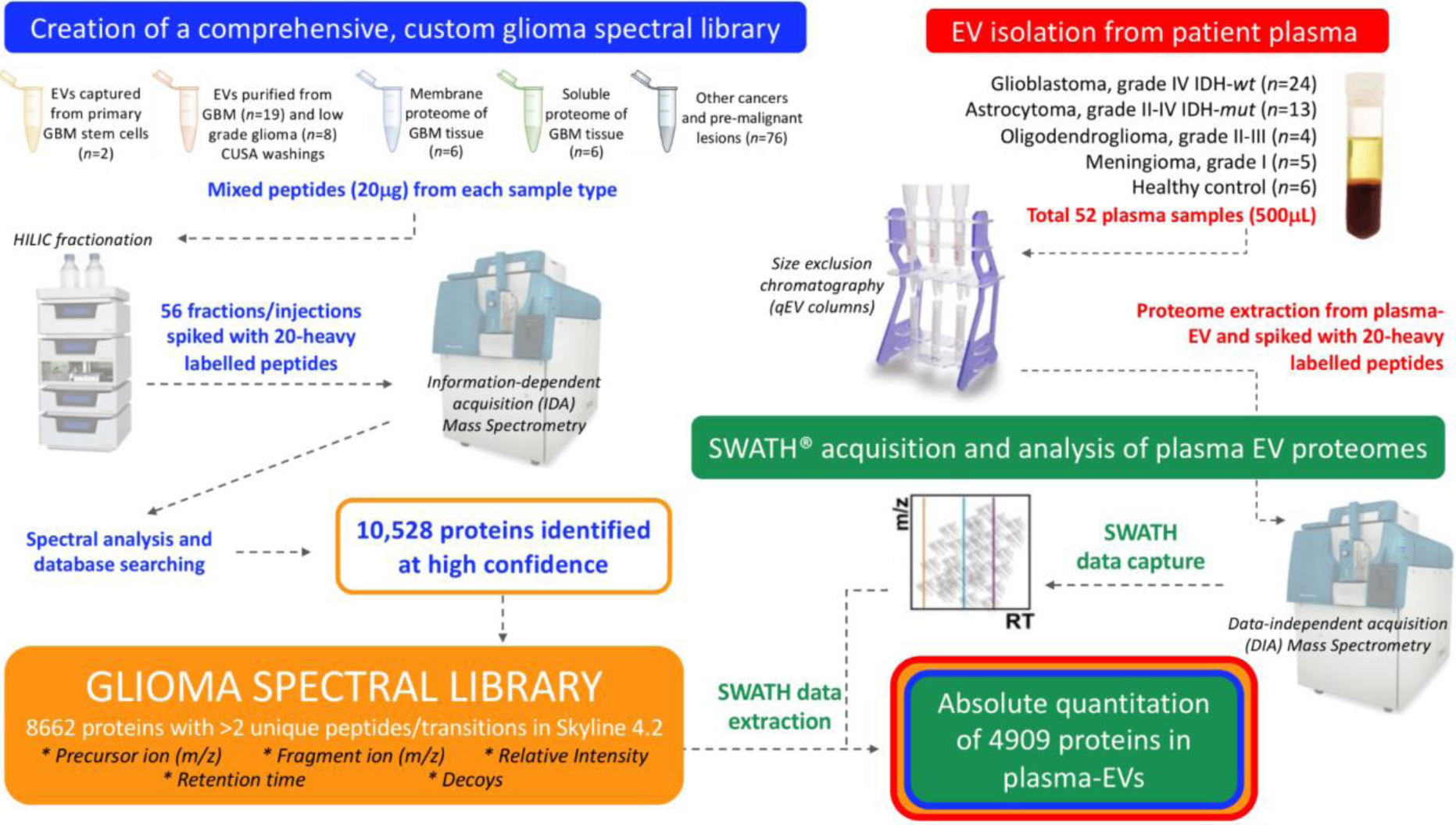
Experimental workflow for in-depth proteomic characterisation of plasma-EVs derived from glioma patients. A custom glioma spectral library was created by information dependent acquisition (IDA)-based LC-MS/MS of hydrophilic interaction liquid chromatography (HILIC) fractionated peptides derived from GBM and other cancer lesions. Database searching of 56 fractions identified 10,528 proteins. Proteins with more than 2 unique peptides and transitions were selected for the creation of a spectral library and comprised of 8662 unique protein species that contained reference sequences (precursor ion (*m/z*), fragment ion (*m/z*), relative intensity, retention time and decoys). Size exclusion chromatography (SEC) isolated plasma-EV proteomes were then assessed by SWATH-MS and the protein data was extracted by aligning the SWATH-MS retention times to the spectral library using 20 heavy-labelled *PepCalMix* peptides that were spiked into both the plasma-EV and spectral library specimens.

The IDA method for analysis of HILIC fractionated peptides involved a MS survey scan range of 350-1700 *m/z* (0.25 sec accumulation time) in positive ionisation mode, with rolling collision energy and dynamic accumulation selected. MS/MS scans were acquired in high sensitivity mode covering a mass range of 100-2000 *m/z* (25 ms accumulation time) of the 20 most intense ions with charge states of 2+ to 5+ and a mass tolerance of 20 ppm. Mass-charge ratios selected for MS/MS were dynamically excluded for 10 s. Prior to loading the samples, a LC-MS/MS standard consisting of 30 fmol pre-digested BSA was injected to test the performance and dynamic range of the instrument. To analyse the plasma-EV peptides, the TripleTOF^®^ was operated in SWATH mode and covered a total of 159 custom variably sized windows (with a 1.0 Da window overlap) over a precursor mass range of 350-1750 *m/z* (Supplementary Table 3, Supplementary Table 1). The IDA and SWATH methods were operated under the same chromatographic conditions on the TripleTOF^®^.

### Creation of a glioma spectral library

IDA LC-MS/MS data captured from HILIC fractionated GBM specimens were added to an established spectral library created from pre-malignant lesions and squamous cell carcinoma [53]. Protein database searching was performed against a reference Human Uniprot FASTA database (03-10-2018) by ProteinPilot™ Software 5.0 (SCIEX; Framingham, MA, USA) using a Paragon™ algorithm. The IDA data was uploaded to ProteinPilot and search parameters were set as follows: *Cys alkylation*-iodoacetamide; *digestion*-trypsin; *instrument*-TripleTOF 6600; *species*-homo sapiens. A detection protein threshold of 10% and false discovery analysis were selected. The resulting spectral library file (.group) was imported to Skyline 4.2 (Maccoss Labs) and label-free analysis of the data was performed. The resultant Protein Pilot spectral library is provided in excel format (Supplementary File A). In Skyline 4.2, only peptides with a confidence score > 0.05 were included in the spectral library for extracting SWATH-MS data and peptide quantitation.

### Targeted Peak Extraction of SWATH-MS Data

Chromatographic peaks were extracted for the plasma-EV samples by aligning the SWATH-MS acquisitions of the plasma-EVs to the comprehensive spectral library in Skyline 4.2, and calibrating the retention times for standard *PepCalMix* peptides. Skyline 4.2 (Maccoss Labs) is an open source software for curating and analysing proteomic data [54] (https://skyline.ms/project/home/software/Skyline/begin.view). Briefly, the spectral library (.group) generated for the IDA files in ProteinPilot™ was imported to Skyline 4.2 and SWATH data files were processed using a full-scan MS/MS resolving power of 30,000. A protein was confidently assigned if it had at least 2 peptides and 2 fragment ions. Peaks were assessed for quality and analysed if at least half of the transitions contributed to a co-eluting peak.

The Skyline document was prepared for SWATH-MS analysis with the peptide and transition settings as illustrated in Supplementary Figure 2 and 3. The human uniprot protein database (3-10-2018) was imported to Skyline and a maximum of 3 tryptic missed cleavages were allowed. Peptides of 5-25 amino acids in length were accepted with variable modifications, oxidation (M) and carbamidomethyl (C). Spectra with precursor charges of 2, 3, 4, ion charges of 1 and 2 for y, b and p ions were filtered for analysis. The MS/MS spectral library was then imported into Skyline, filtering spectra that passed a q-value cut-off score of 0.95, and removing peptides that a) matched multiple proteins, b) peptides that did not match any proteins, or c) peptides that did not meet the filter settings. Before importing the SWATH-MS data to Skyline, an isolation scheme was created to inform Skyline of the 159 variable isolation windows that had been used to acquire the SWATH-MS data on the TripleTOF^®^ 6600 (Supplementary Figure 4). The raw SWATH-MS (.wiff) acquisitions of plasma-EV samples were imported to Skyline and reintegration of the peaks was performed to discriminate between the target peaks and decoys. Peaks with a q-value < 0.05 were reintegrated. To allow more accurate retention time (RT) prediction, an indexed retention time (iRT) calculator for *PepCalMix* peptides (Supplementary Figure 5A) was calibrated to the RT values for the iRT peptides that were measured in the spectral library. SWATH-MS scans were used if their RT was within a 5 min window of the predicted RT. MSstats input files were generated to acquire values for all identified proteins.

### Differential expression analysis and statistics

Following peak identification and quantitation, group comparisons were performed for identified proteins with MSstats3.7.3 (http://msstats.org/), an open source R package in Bioconductor [55]. Before MSstats could be used for differential expression (DE) analyses, biological replicates were annotated into their respective conditions and MSstats Input reports were generated in Skyline4.2 and analysed in RStudio 1.0.153 and R3.5.0. MSstats input reports for SWATH experiments contained variables of protein name, peptide sequence, precursor charge, fragment ion, product charge, isotope label type, condition, bioreplicates, run and intensity. Firstly, MSstats3.7.3 pre-processed the input reports to the right format by removing iRT proteins, allocating NAs for truncated peaks and assigning 0 for intensities with a detection q-value less than 0.01. The data was then processed using the default MSstats settings which involved log_2_ transformation, normalisation by equalising the median intensities across runs, model-based quantification by Tukey’s median polish (TMP) and imputation of censored missing values using the accelerated failure time (AFT) model. Group comparisons were performed to yield fold change (FC), log_2_ fold change (log_2_FC), standard error of the log_2_ fold change (SElog_2_FC), student t-test *p*-value, and Benjamini-Hochberg corrected *p*-values with a 5% FDR threshold.

Principal component analysis (PCA) was performed on samples based on protein expression to visualise sample clustering. The top-50 DE significant proteins across all comparisons were identified by filtering and ranking proteins by their adjusted *p*-value and FC. Significant proteins were visualised with heat-maps plotted in the R environment. Heat-maps of the top-50 significant proteins were plotted as log_2_-transformed FC, calculated by subtracting the protein abundance from its mean value for that protein across all samples. Pathway analysis was performed using Ingenuity^®^ software (Ingenuity Systems, USA; http://analysis.ingenuity.com) to assess functional associations (biological and canonical pathways) of DE proteins between the plasma-EVs of GBM and healthy controls by performing core expression analyses using default criteria. The PANTHER over-representation test (http://www.pantherdb.org) was used to perform gene ontology enrichment analysis of biological functions and reactome pathways for the DE proteins across the genetic-histological glioma subtypes and controls. The over-representation test was performed against the *Homo sapiens* reference database and a Fisher’s Exact Test was performed and corrected at a 5% FDR.

### Data availability

The custom IDA glioma-spectral library and raw SWATH-MS acquisitions of the plasma-EV peptides have been deposited to ProteinAtlas with an identifier number: PASS01487 (www.peptideatlas.org/repository).

## Results

### Characterisation of plasma-EVs

EV elution and size distribution profiles for all twelve plasma-EV fractions were consistent across plasma sampled from three healthy individuals. A discernible EV population was first observed to elute in fraction 7, and the EV concentration increased for each subsequent fraction (Figure 2A). EV containing fractions 7-12 were predominantly comprised of small vesicle subtypes of less than 200 nm, with modal size distributions of 80-100 nm and a vesicular morphology (Figure 2B, C). Investigating the proteomes of the EV containing fractions 7-12 by quantitative label-free LC-MS/MS determined the fractions most suitable for blood-based biomarker discovery, i.e. fractions with the highest coverage of EV-related proteins and lowest expression of soluble blood proteins. The highest number of unique protein IDs were detectable by LC-MS/MS in fractions 7-9 and corresponded to more confident EV-related protein identifications (Figure 2D). Functional enrichment analysis found that plasma-EV fractions 7-12 had significant annotations to exosomes and other membranous compartments (*p*<0.01; Figure 2E), with higher annotations for fractions 7-9 compared to 10-12 (Figure 2E). Overall, a total of 82 of the top-100 canonical EV proteins, as reported by ExoCarta, were identified in the plasma-EV fractions and were detected across fractions 7, 8 and 9 (Supplementary File B). The higher number of unique protein identifications by LC-MS/MS in the earlier eluting EV fractions coincided with lower expression of common highly-abundant serum-related proteins (Figure 2F), which increased with each sequential EV fraction (Figure 2F); there were increases in A2M (5.3-fold), C3 (3.4-fold), TF (3.3-fold), CFH (23.4-fold), HP (5.3-fold) and IGG1 (2.7-fold) in fraction 12, relative to fraction 7. As such, the earlier eluting plasma-EV fractions 7-9 were considered the most attractive candidates for large-scale proteomic analysis for EV biomarker discovery.

**Figure 2.**
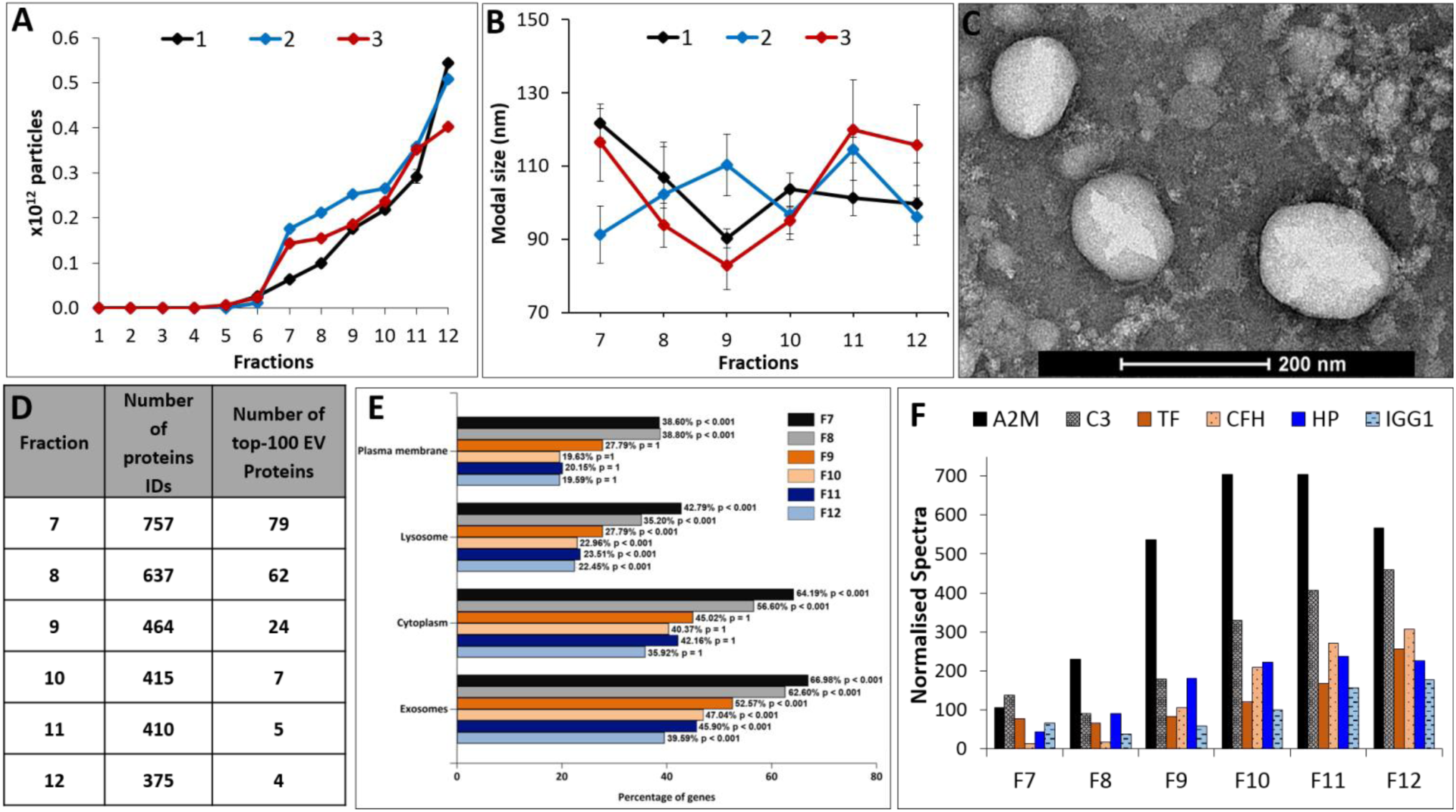
Characterisation and annotations of plasma-EV fractions. Nanoparticle tracking analysis (NTA) determined (A) number of particles and (B) modal size distribution for twelve 500 µL PBS EV fractions (F1-12) isolated from the plasma of three healthy individuals (1, 2, 3). The results are shown as the mean ± standard error of 3 independent NTA measurements. (C) Close-up TEM image of F7-9 EVs, 200 nm scale bar. (D) The number of confident protein identifications, including the top-100 EV-related proteins as reported by ExoCarta, determined by LC-MS/MS measurements of F7-12 EVs. (E) Functional enrichment annotations to cellular compartments for the LC-MS/MS sequenced F7-12 EVs. (F) Quantitative LC-MS/MS measurements of highly-abundant serum proteins problematic for biomarker discovery of complex biological specimens across plasma-EV fractions 7-12.

### Assembling a spectral library tailored to glioma SWATH analysis

IDA LC-MS/MS analysis of HILIC fractionated peptides derived from glioma and other tumour specimens included a total of 10,528 protein species identified at an unused ProtScore detection threshold of 0.05. Of these proteins, 9414 had a maximum global FDR of 5% and were included in the spectral library. Of these proteins, 8662 proteins in the library were most optimal for targeted proteomics as they had at least two unique peptides and transitions, and were selected to be included in the Skyline target list for SWATH data extraction (Supplementary File A).

### SWATH-MS data extraction, quality and reproducibility

Plasma-EV proteomes (*n*=52 total samples; 41 glioma II-IV, 5 non-glioma controls and 6 healthy controls; Table 1), were sequenced by SWATH-MS and the peaks were aligned to the spectral library. To ensure accurate assignment of the SWATH-MS peaks, the retention times of the SWATH-MS peaks were aligned to the IDA spectral library using an indexed retention time (iRT) calculator generated from twenty heavy-labelled *PepCalMix* reference peptides spiked and measured in both plasma-EV peptides and spectral library specimens. The iRT linear regression had a Pearson’s correlation coefficient (*r*^*2*^) of 0.9903, reflecting highly correlative measured retention times and *PepCalMix* iRT definition values (Supplementary Figure 5A). The retention times of the *PepCalMix* peptides were also highly reproducible across all plasma-EV samples (Supplementary Figure 5B), and normalised peak areas of the standard peptides were similar across the sample groups (Supplementary Figure 5C). This ensured that the plasma-EV peptides analysed by SWATH-MS could be reliably and reproducibly aligned to the spectral library for accurate quantitative comparisons. The SWATH-MS data quality was further assessed by manual inspection of randomly selected extracted ion chromatographs (XIC) for *PepCalMix* peptide, SPYVITGPGVVEYK, and Aurora Kinase A peptide, VLCPSNSSQR to assess peak shape, retention time stability, signal intensity and signal-to-noise ratio (Supplementary Figure 6). All parameters were highly reproducible across samples, with identical peak shapes observed that had minimal noise and less than 1 min chromatographic retention-time shifts across samples.

### SWATH-MS based characterisation of glioma plasma-EV proteomes

A total of 4909 plasma-EV proteins with q-values ≤ 0.05 were confidently aligned to the IDA spectral library for all 52 plasma samples. Grouped by genetic-histological subtype, the MSstats package filtered the aligned proteins and confidently identified 4834 proteins in *IDH-*wt GBM (GBM), 4814 proteins in *IDH-*mut astrocytoma (AST), 4270 in *IDH-*mut oligodendroglioma (OLI), 4488 in benign grade I meningioma (MEN), and 4586 in healthy control (HC) plasma-EVs with q-values≤0.01. Similarly, when grouped by glioma grade II-IV (GII-IV), 4836 proteins were identified for GIV, 4750 in GIII and 4730 in GII with q≤0.01 (Supplementary File C). FunRich analysis of the plasma-EV proteins found that all the glioma subtypes had similarly significant (p<0.01) functional annotations. A significant percentage of the plasma-EV proteins were mapped to biological processes such as cell growth and/or maintenance (7.9 ± 0.07 %), transport (9.17 ± 0.08 %) and metabolism (14.79 ± 0.15 %) across the glioma genetic-histological subtypes and controls. Interestingly, the plasma-EV proteins also had significant association with sites of expression such as the brain (64.87 ± 0.35 %) and malignant glioma (53.98 ± 0.33 %). MSstats confidently identified 4054 proteins that were common to plasma-EVs from all sample cohorts, 90% of which were annotated as ‘extracellular vesicle’, ‘exosome’, ‘microparticle’ and ‘microvesicle’ proteins in Vesiclepedia (Figure 3A). An unsupervised cluster analysis by principal component analysis (PCA) of the samples based on plasma-EV protein expression found a high level of discrimination between EVs sampled from the plasma of patients with highly malignant gliomas (GBM/GIV) and controls (HC and MEN) (Figure 3B-1, B-2), while less malignant glioma phenotypes (GII-III, Figure 3B-1; AST and OLI, Figure 3B-2) were intermediately dispersed.

**Figure 3.**
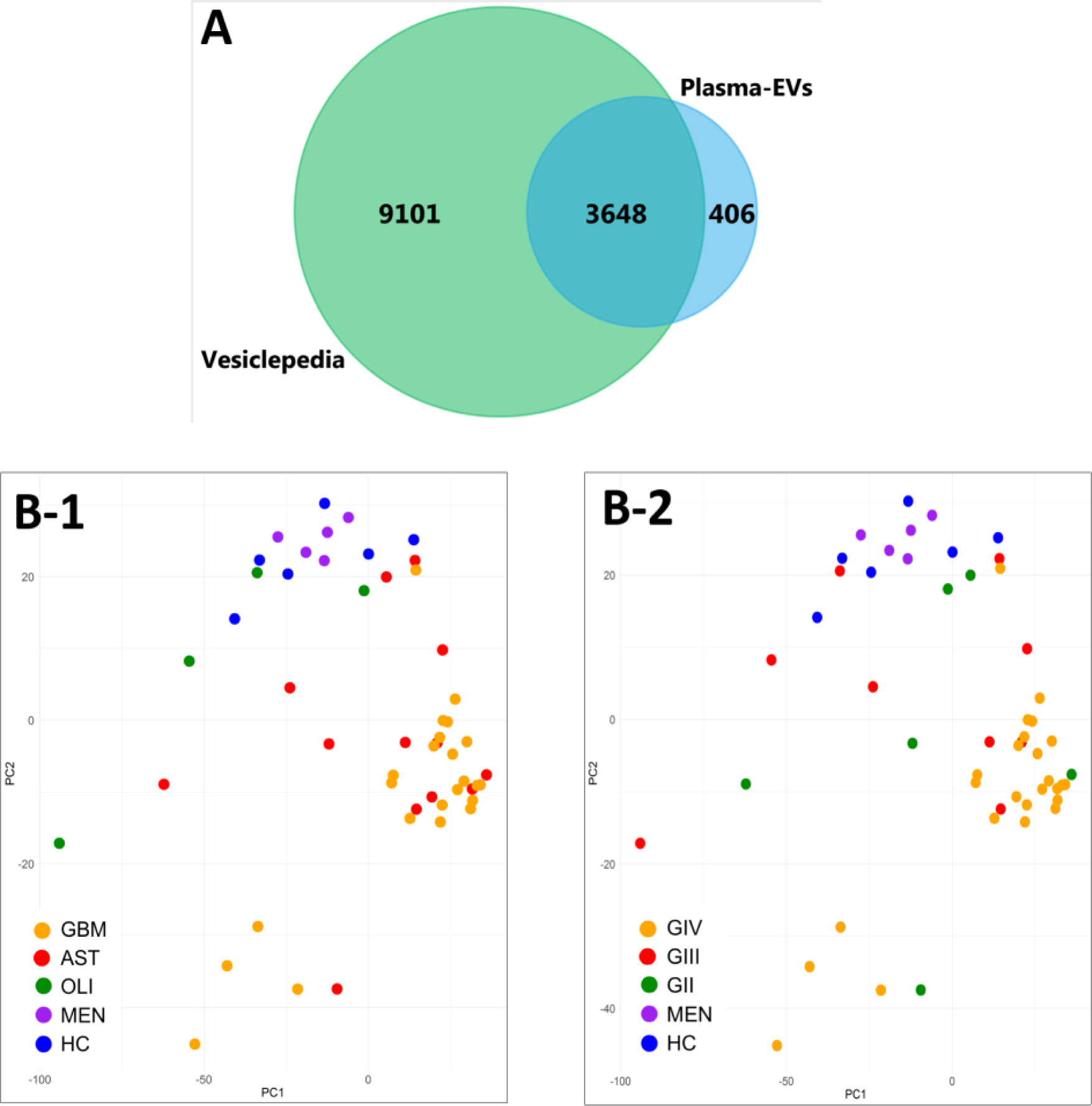
Characterisation of plasma-EV proteins by SWATH-MS. (A) Venn diagram showing overlap between the 4054 proteins confidently identified in the plasma-EVs in all sample groups by MSstats (blue) to EV proteins compiled in Vesiclepedia (green; matched by gene name; restricted to proteins annotated as originating from “extracellular vesicles”, “exosomes”, “microparticles” and “microvesicles”). (B) Unsupervised clustering by principal component analysis (PCA) based on the plasma-EV protein expression for samples grouped by their respective (B-1) genetic-histological subtype; GBM (orange), AST (red), OLI (green), MEN (purple) and HC (blue); the proportion of the variance is explained by PC1 (x-axis) = 18.48% and PC2 (y-axis) = 7.28% (y-axis), and by (B-2) glioma grade; GIV (orange), GIII (red), GII (green), MEN (purple) and HC (blue); the proportion of the variance is explained by PC1 (x-axis) = 18.48% and PC2 (y-axis) = 7.28% (y-axis).

### Significant proteins in glioma plasma-EVs

A total of 463 plasma-EV proteins were identified as significantly DE when samples were grouped according to the glioma genetic-histological subtypes (adj. *p*-val≤0.05; Supplementary File D). Similarly, 318 proteins significantly changed in plasma-EV samples when samples were grouped by glioma grade (adj. *p*-val≤0.05; Supplementary File D). Gene ontology enrichment by PANTHER revealed that the signficant plasma-EV proteins across glioma subtypes were predominantly associated with energy generation by mediating proton transport (6.06-fold enrichment, *p*=0.0491), vesicle-mediated transport (2.12-fold enrichment, *p*=0.022), carboxylic acid biosynthesis (4.99-fold enrichment, *p* = 0.0194) and protein folding (3.88-fold enrichment, *p*=0.0352). Signifcant enrichment in the plasma-EV proteins was implicated to reactome pathways such as the WNT5A-dependent internalisation of FZD4 (12.12-fold enrichment, *p*=0.0435), EPH-Ephrin signaling (7.76-fold enrichment, *p* = 0.011), innate immunity (2.47-fold enrichment, *p* = 1.6×10^−7^) and vesicle-mediated transport (3.03-fold enrichment, *p*=2.94×10^−8^) (Supplementary Figure 7B). Functional pathway analysis by Ingenuity^®^ for DE proteins between GBM and HC plasma-EVs (463 proteins; FC≥2, adj. *p-val*≤0.05) revealed significant associations to cancer (*p-val* range: 2.28×10^−2^–8.10×10^−19^, 450 molecules) and clathrin-mediated endocytosis signalling (*p*=1.37×10^−4^). The top scoring molecular and cellular functions included cellular function and maintenance (*p-val* range: 2.07×10^−2^–8.67×10^−9^, 104 molecules), protein sythesis (*p-val* range: 2.05×10^−2^–6.42×10^−5^, 80 molecules) and post-translational modification (*p-val* range: 2.28×10^−2^–3.19×10^−4^, 24 molecules) (Supplementary Figure 7A).

Distributions of the top-50 DE proteins across the genetic-histological subtypes and WHO grades II-IV are displayed as heat-maps (Figure 4A). PCA visualisation of sample clustering based on the expression of the top-50 proteins revealed a tendency for glioma samples to group away from controls and with their respective genetic-histological subtype (Figure 4 B-1) or grade (Figure 4 B-2). The samples derived from glioma patients with more aggressive phenotypes (GBM/GIV) were highly distinguishable from controls, clustering to the far left of the PCA, while plasma-EVs derived from less aggressive gliomas (AST and OLI, Figure 4B-1; GII-III, Figure 4B-2) had a less defined cluster and were scattered between the highly aggressive GBMs and controls. Interestingly, plasma-EV profiles collected from three patients with recurrent progressions of their tumours, shifted towards more aggressive glioma phenotypes. The EV profile of a grade III *IDH-*mut astrocytoma patient that progressed from grade II (*circle*: AST, Figure 4C; GII to GIII, Figure 4D) shifted to the left of the PCA, closer to the grade IV GBM samples. A grade IV *IDH-*mut astrocytoma blood sample that had progressed from a prior grade III (*star*: AST, Figure 4C; GIII to GIV, Figure 4D) also shifted to the left and banded with the majority of grade IV GBM tumours. In addition, a recurrent sample derived from a patient with an *IDH-*wt GBM (*rectangle*: GBM, Figure 4C; GIV, Figure 4D) also had an observable shift to the left of the PCA, conforming to a pattern of more aggressive disease.

**Figure 4.**
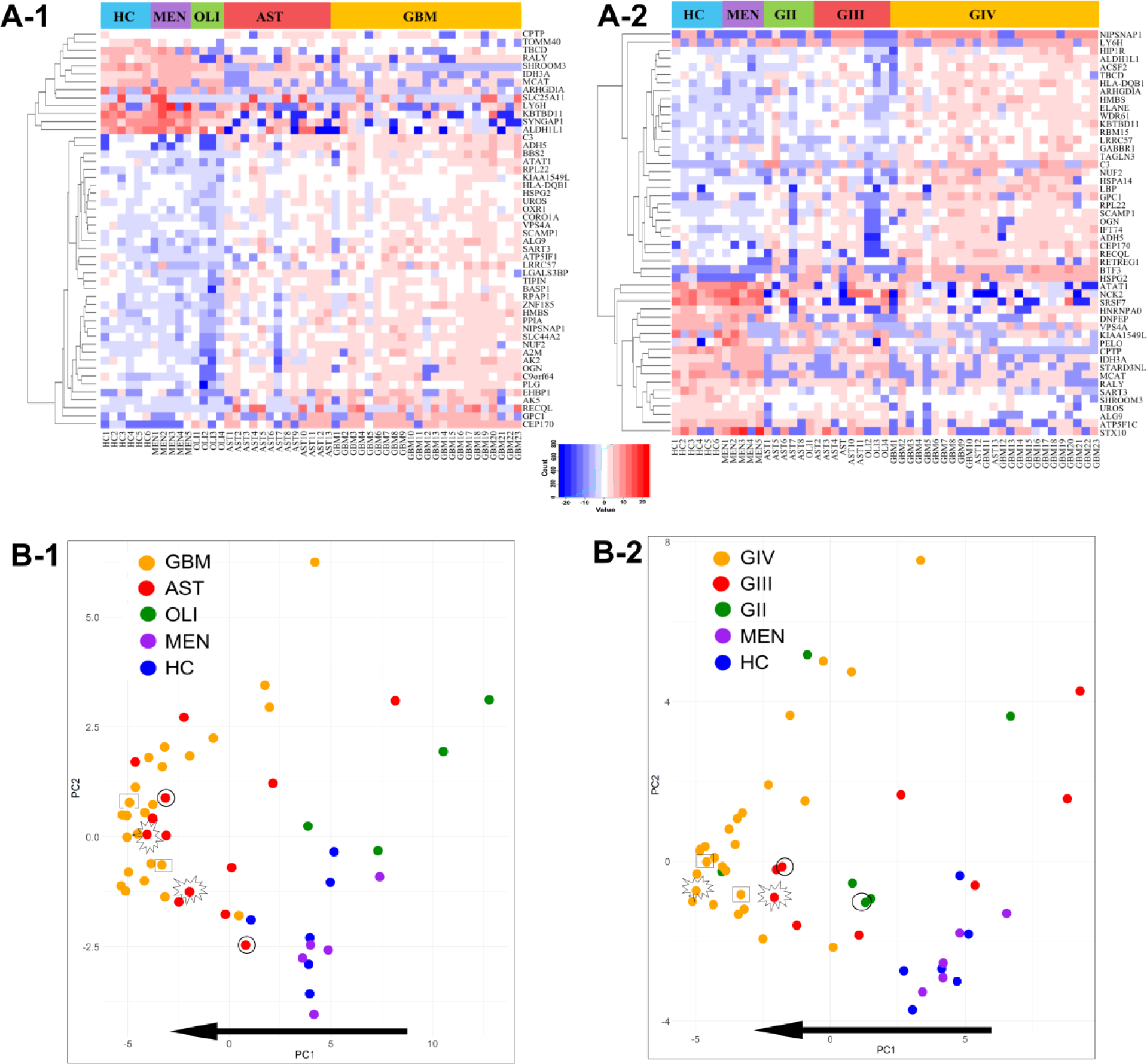
Visualisation of the top-50 differentially-expressed (DE) proteins identified across glioma plasma-EVs. Heat-map representations of the top 50 significant DE plasma-EV proteins for samples categorised by (A-1) genetic-histological subtype and (A-2) grade. Principal component analysis (PCA) based on the expression of the top-50 plasma-EV proteins by their respective (B-1) genetic histological subtype; HC (blue), MEN (purple), OLI (green), AST (red) and GBM (orange); the proportion of the variance is explained by PC1 (x-axis) = 42.95% and PC2 (y-axis) = 8.36% and (B-2) grade; HC (blue), MEN (purple), GII (green), GIII (red) and GIV (orange); the proportion of the variance is explained by PC1 (x-axis) = 32.71% and PC2 (y-axis) = 11.49%. (B-1, B-2) Glioma samples that are circled were derived from an AST patient that progressed from GII to GIII. Samples that are starred were derived from an AST patient that progressed from GIII to GIV, while samples that are enclosed in a rectangle were derived from a recurrent GBM (GIV). The arrow shows the direction of increased glioma aggression.

### Putative plasma-EV biomarkers for distinguishing glioma subtypes

The plasma-EV proteomes were interrogated to identify proteins with restricted expression in the glioma subtypes. A total of 11 proteins were only detected in plasma-EVs from GBM patients, i.e., AIDA, ARHGEF10, BNIP3L, FYB1, KMT2D, MAP7, MAST4, PDE8A, POLR2D, RENBP and SLC25A17 (Figure 5A). The expression of CDC40 was exclusive to AST plasma-EVs and TPST2 restricted to OLI plasma-EVs. There were no unique plasma-EV proteins in control specimens. Correspondingly, the 11 GBM plasma-EV proteins were also restricted to GIV samples, in addition to CETN3, PPP1R11 and SYT7. The CDC40 and TPST markers detected in AST and OLI samples respectively, showed restricted expression in GIII patients, while no proteins were specific to GII patients (Figure 5A). The relevance of the restricted proteins in GBM/GIV plasma-EVs was further investigated *in silico* to determine whether their expression was a result of EV selective packaging or if they were reflective of molecular changes associated with GBM pathology. The transcript levels corresponding to the GBM/GIV restricted proteins were generated by Oncomine (Compendia Biosciences, MI, USA) using TCGA data (GBM, *n*=542; normal brain, *n*=10; Human Genome U133A Array, TCGA 2013 [56]). Significant differences in transcript levels were observed between GBM and normal brain tissue for AIDA (2.5-fold, *p*=3.4×10^−8^), BNIP3L (1.2-fold, *p*=5.5×10^−6^), CETN3 (1.8-fold, *p*=1.7×10^−7^), FYB1 (1.8-fold, *p*=7.25 ×10^−9^) and POLR2D (1.7-fold, *p*=8.14 ×10^−5^).

**Figure 5.**
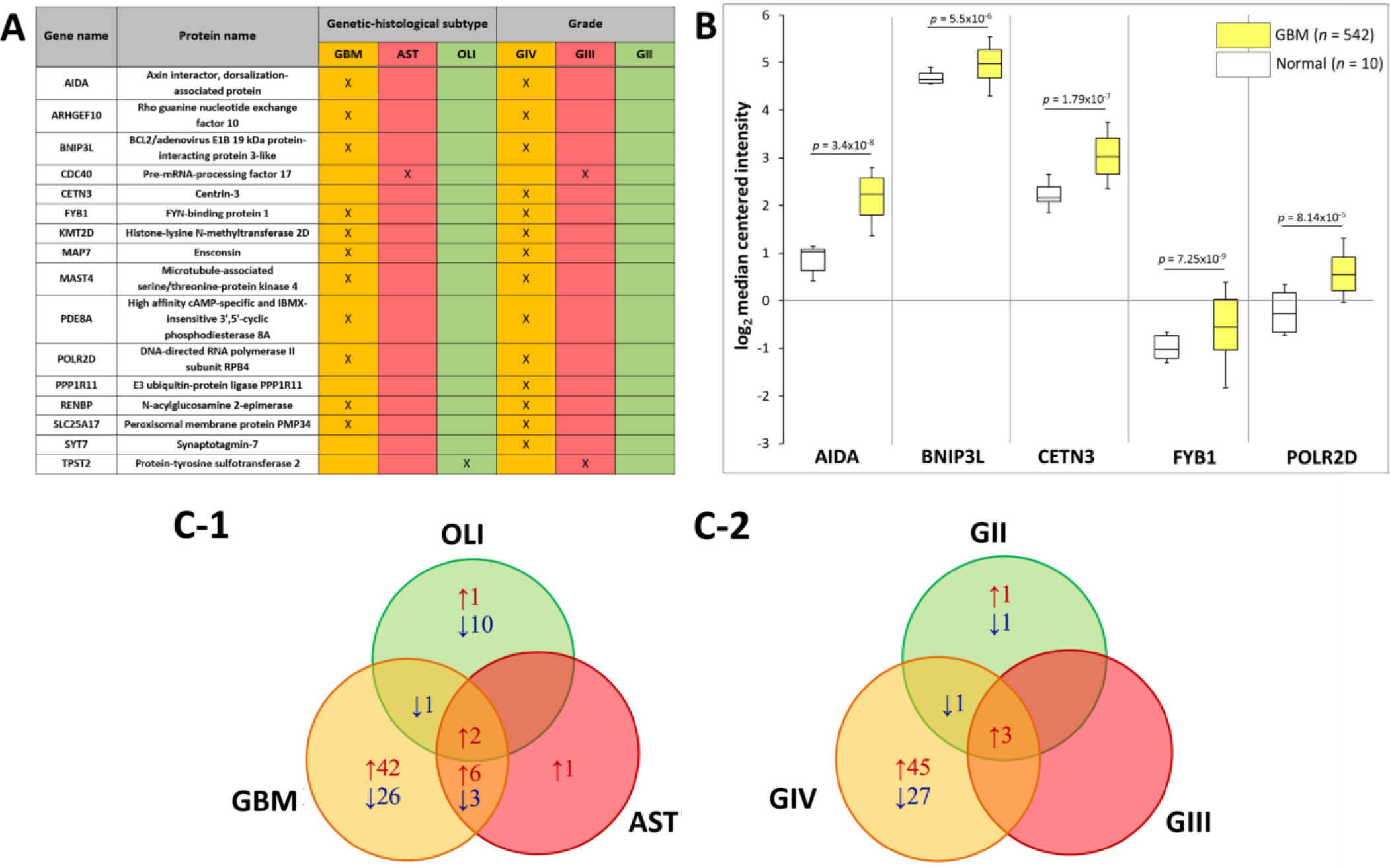
Putative plasma-EV protein markers for glioma. (A) Proteins with restricted expression in plasma-EVs according to different glioma genetic-histological subtypes (GBM, AST, OLI) and WHO grades (GIV, GIII, GII). Proteins that are present in the plasma-EVs of their respective glioma subtype are denoted by X. (B) Box-plots were generated by Oncomine for plasma-EV proteins restricted to GBM/GIV patients that revealed significant relative GBM gene expression levels (represented by log2 median centred intensity; U133A Array, TCGA 2013). Significant transcripts in GBM tissue included AIDA (2.5-fold, *p*=3.4×10^−8^), BNIP3L (1.2-fold, *p*=5.5×10^−6^), CETN3 (1.8-fold, *p*=1.7×10^−7^), FYB1 (1.8-fold, *p*=7.25 ×10^−9^) and POLR2D (1.7-fold, *p*=8.14 ×10^−5^). *n* is the number of samples, the 90^th^ percentile is signified by upper error bars, and the 10^th^ percentile is represented by the lower error bars. (C) Plasma-EV protein changes that were significant in glioma relative to both healthy and non-glioma controls (*p*-val≤0.05, |fold-change|≥2). The Venn diagrams show overlap of the glioma-related plasma-EV proteins in (C-1) GBM, AST and OLI and (C-2) GIV, GIII and GII plasma-EVs, where arrows denote direction of fold change. The identities of glioma-related plasma-EV proteins are listed in Supplementary Tables 4 and 5.

DE analysis comparing GBM, AST and OLI plasma-EVs to HCs, revealed 184, 33 and 18 significantly changing proteins, respectively (adj. *p*-val≤0.05, FC≥2). Similar patterns were observed relative to MEN, with 179, 45 and 22 DE proteins in GBM, AST and OLI, respectively (adj. *p*-val≤0.05, FC≥2). Similarly, GIV, GIII and GII glioma samples had 178, 6 and 12 proteins with significantly different expression relative to HC, respectively, while 182, 19 and 17 proteins were DE in GIV, GIII and GII, relative to MEN, respectively (adj. *p*-val≤0.05, FC≥2). The complete list of significantly changing proteins is provided in Supplementary File D. To further assess whether EV proteomes are able to distinguish glioma from non-glioma plasma, proteins with significant expression relative to both control groups (HC and MEN) were collated. The glioma-associated proteins included 80, 12 and 14 proteins that were DE in GBM, AST and OLI plasma-EVs, relative to both controls. The overlap of the proteins across the glioma subtypes and the direction of change relative to controls are displayed as Venn diagrams (Figure 5C-1) and listed in Supplementary Table 4. Two proteins, EBNA1BP2 and FAM129A, were significantly expressed in GBM, AST and OLI samples relative to the controls. Similarly, when grouped by WHO grade, 78, 3, and 8 proteins were significant in GIV, GIII and GII samples relative to both controls, respectively. The overlap of grade associated plasma-EV protein changes and the direction of change relative to controls is displayed as a Venn diagram (Figure 5C-2) and the proteins are listed in Supplementary Table 5. Three proteins showed differential abundance in all grades (II-IV) relative to the controls, AK2, CYB5A and GOLT1B.

### Overlap of significant plasma-EV proteins with previous GBM-EV findings

SWATH-MS analysis of glioma plasma-EVs allowed the identification of proteins already associated with GBM-EVs. The plasma-EV SWATH-MS data was mined for the previously reported *in vitro* GBM-EV signature proteins, which comprised of 145 proteins shared by EVs derived from six established GBM cell-lines [25]. A 90% overlap was observed between the glioma plasma-EVs and the *in vitro* protein signature, with 130 proteins measured in plasma-EVs of glioma patients. Of these, 10 ‘signature’ GBM-EV proteins [25] were significantly expressed in the plasma of GBM patients relative to either HC or MEN, including PSAP, CALR, PLOD3, HSPA4, GANAB, LGALS3BP, CCT2, PPIA, C3 and KRT10 (Figure 6A). Proteins previously identified to be significantly expressed in EVs captured from neurosurgical aspirates of GBM compared to grade II-III glioma (298 proteins) [24] were also identified in EVs captured from plasma with 87% (260 proteins) overlap. Twelve neurosurgical aspirate EV proteins also showed significance in GBM plasma-EVs relative to HC or MEN, including ANXA2, UQCRC2, COX5A, NDUFS4, IARS, TMEM65, CCT7, PSMC3, CCT2, GABBR1, SCD5 and TOMM40 (Figure 6A). Previously identified GBM-EV proteins [24, 25], PSAP (Figure 6B), PPIA (Figure 6C), CCT7 (Figure 6D) and C3 (Figure 6E) also changed significantly in the plasma-EVs of GBM/GIV patients relative to both HC and MEN (Supplementary File D).

**Figure 6.**
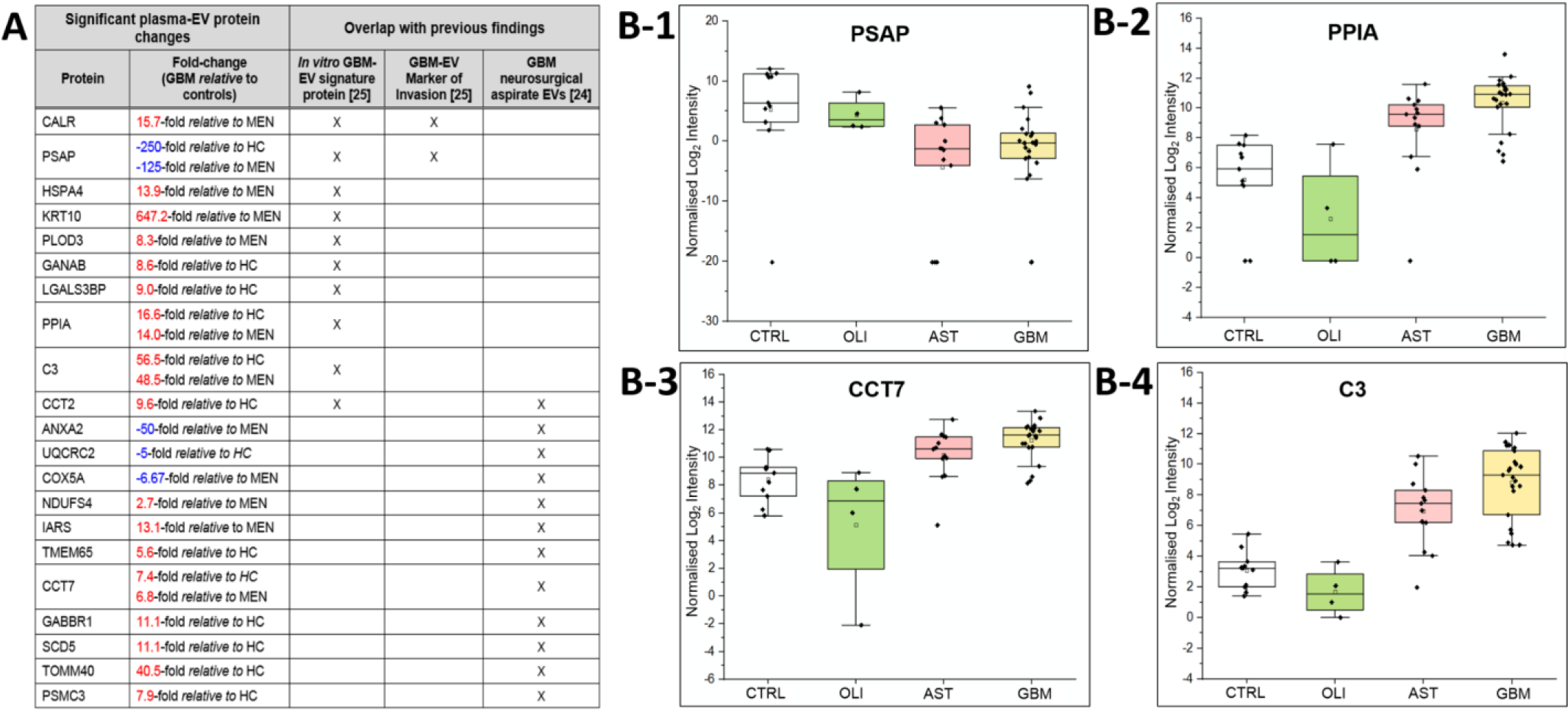
Overlap of significant GBM/GIV plasma-EV proteins with previous GBM-EV findings. (A) Proteins that are significant in GBM plasma-EVs relative to either MEN or HC (|fold-change| ≥ 2, adjusted *p-val* ≤ 0.05) previously reported as *in vitro* GBM-EV signature proteins [25] or to have significance in EVs derived from GBM neurosurgical aspirates [24]. Significant plasma-EV proteins that overlap with previous GBM-EV findings are denoted by X. (B) Box-plots show the distribution of significant plasma-EV proteins, (B-1) PSAP, (B-2) PPIA, (B-3) CCT7 and (B-4) C3. The box-plots depict the expression levels for CTRLs (total controls, *n*=11; non-glioma, *n*=5 and healthy controls, *n*=6; white), OLI (*n*=4; green), AST (*n*=13; pink) and GBM (*n*=23; yellow). The protein expression for each individual patient is plotted as normalised log2 intensity and denoted by a black dot. The upper error bars signify the 90^th^ percentile, and lower error bars represent the 10^th^ percentile, the middle line represents the median and open square signifies the mean.

## Discussion

Accurate and sensitive glioma blood-based biomarkers will arm clinicians with much-needed information and offer patients more definitive and timely answers about the active state of their cancer and the effectiveness of their treatment. Circulating-EVs hold real promise as robust and readily accessible pools of glioma biomarkers. While high-throughput next-generation-sequencing technologies have allowed thorough characterisation of nucleic acids in blood-derived EVs [30, 57], the proteome of circulating-EVs remains largely unexplored. Shot-gun proteomic methods have been primarily used to investigate glioma-EV proteomes [58, 59]. However, such approaches achieve limited proteomic coverage, especially for complex blood-derived specimens [60]. We have employed SWATH-MS in conjunction with a data extraction strategy against a comprehensive spectral library for in-depth assessment of the proteomic content of circulating-EVs from glioma patients. The expansive coverage of targeted SWATH-MS allowed the identification of potential peripheral EV-associated glioma biomarkers, as well as the translation of previous *in vitro* [25] and *ex vivo* findings [24].

In the literature, there are no definitive propositions for the use of plasma over sera for EV biomarker discovery. The process to separate serum from blood first requires blood coagulation, which causes platelet activation and leads to increased platelet-EV secretion [61]. Sera also comprises a higher proportion of particles larger than 200 nm, compared to plasma [62] and thus plasma was selected as the starting material to optimise our biomarker workflow. SEC was chosen to isolate plasma-EVs for discovery proteomic analysis as it is rapid and reliable for isolation of small-EV subtypes from complex biological fluids. While SEC retains soluble protein and lipoprotein contaminants in EV preparations [64, 65], it is readily adaptable, scalable and automatable, making it ideal for routine EV isolation in diagnostic pathology service environments. The suitability of our discovery plasma-EV proteomics approach was evaluated using SEC fractions from healthy plasma (*n*=3) and the isolated EV populations were characterised to meet the minimal information for studies of extracellular vesicles 2018 (MISEV 2018) criteria [66]. The presence of canonical EV proteins and serum protein contaminants (Figure 2) determined that plasma-EV fractions 7-9 were the most suitable for large-scale proteomic analysis for EV biomarker discovery.

### SWATH produces high-quality proteomic data for plasma-EV biomarker discovery

A comprehensive custom 8662-protein spectral library was constructed using fractionated peptides derived from multiple glioma specimens and other cancer-related material to allow for the targeted identification and quantitation of plasma-EV proteins. Circulating-EV peptides from diffuse glioma patients and controls were prepared and analysed by SWATH-MS. A targeted extraction method was used to align SWATH-MS retention times to the custom spectral library. The retention-time alignment was facilitated by *PepCalMix* standard peptides, spiked into both the library and plasma-EV specimens. Using a mProphet scoring algorithm and FDR estimation, this proteomic workflow facilitates confident and reliable peptide quantification across the plasma-EV biological replicates [67], with highly reproducible retention-times and XIC peptide peaks across all plasma-EV samples (Supplementary Figure 5A-C, Figure 6). Overall, we achieved a total of 4054 high-confidence protein identifications across the plasma-EV sample groups; the identified proteins had significant functional associations to exosomes and a high degree of overlap with Vesiclepedia-listed EV proteins (Figure 3A). These results demonstrate the capability of state-of-the-art SWATH-MS technology for circulating-EV proteome characterisation, allowing in-depth proteomic assessments of complex specimens while maintaining a high level of precision, reproducibility and accuracy across samples.

### Plasma-EV proteomes reflect glioma aggression profiles

The value of SWATH-MS as a biomarker discovery platform was further demonstrated by the observed clustering of glioma subtypes based on plasma-EV protein expression levels, with plasma-EV profiles from diffuse glioma patients readily discernible from healthy and non-glioma controls (Figure 3B, Figure 4B). Moreover, a ‘glioma aggression trend’ was observed, where samples derived from patients with the most aggressive subtypes (GBM, GIV) clustered the furthest away from the controls, with less malignant subtypes (AST, OLI or GII-III) dispersed between the GBM/GIV and control groups. Interestingly, the AST and OLI patient samples were separated on the PCA plots. OLI carries a better prognosis than AST [68], and OLI plasma-EV profiles were distinguishable from AST, aggregating in closer proximity to the control groups, whereas AST clustered towards the GBM samples (Figure 3B-1, Figure 4B-1).

Intriguingly, resampled plasma specimens from patients with confirmed tumour progression exhibited plasma-EV profiles that shifted towards the GBM/GIV cluster, corresponding to more ‘aggressive’ disease (Figure 4B-1, B-2). This observation highlights the potential of sampling blood-EVs as a bona fide approach to monitor glioma patients for recurrent progression. While *IDH*-mut status can indicate glioma severity, grade IV GBMs were not readily distinguishable on the PCA plots by *IDH*-mut status (ie. GIV *IDH*-mut AST versus *IDH*-wt GBM). There were limited specimens from patients with secondary GBM (GIV IDH-*mut* AST) studied; thus, significant plasma-EV proteins between the *IDH*-wt GBM and *IDH*-mut AST categories most likely reflect differences in WHO grade (i.e., GIV vs GII-III). Four proteins were identified with significantly lower levels in *IDH*-wt GBM relative to *IDH*-mut AST, i.e., TIPIN (−50-fold), NUF2 (−16.7-fold), SMARCE1 (−10-fold) and PIEZO1 (−2.4-fold; Supplementary File D), and the utility of these discriminating protein markers should be further investigated.

Interestingly, AIDA, BNIP3L, CETN3, FYB1 and POLR2D were found to be restricted to plasma-EV cargo of GBM patients. Significant expression was also reciprocated for these markers in GBM tissue compared to normal brain in TCGA cohorts (Figure 5B). Furthermore, the open-access resource, the Human Pathology Atlas as part of the Human Protein Atlas (www.proteinatlas.org/pathology), reports high AIDA, CETN3, FYB1 and POLR2D expression to be unfavourable prognostic markers across multiple cancer types [69]. Overall, these results indicate that plasma-EV proteins have significant potential to distinguish patients with more aggressive diffuse glioma, which may be expected, given that higher-grade lesions are more infiltrative and have higher ki-67 proliferative indices [70] and as such, likely secrete larger quantities of tumour-derived EVs containing more oncogenic protein species into the circulation [25, 71, 72].

### Links to previous GBM-EV studies

Multiple protein species resolved in GBM/GIV plasma-EVs here were previously identified in GBM-EV *in vitro* [25] and *ex vivo* [24] studies (Figure 6A). Notably, PPIA, PSAP, CCT7 and C3 were significantly DE in GBM plasma-EVs, relative to both healthy and non-glioma control (Figure 6B; Supplementary File D). PPIA, also known as cyclophilin A, has roles across a range of biological processes, with essential functions in protein folding and chaperone activity [73]. PPIA overexpression is reported in various cancer types, its expression is influenced by chemotherapy [74, 75] and PPIA has been reported to be a target gene of dysregulated miRNA species that are present in the plasma of GBM patients [76]. C3 is a member of the human complement system with key functions in innate immunity. C3 deposits in GBM tissue, suggesting a role of the complement cascade in GBM pathogenesis [77]. PSAP is a highly conserved glycoprotein and precursor of lysosomal proteins (saposins A-D) that plays substantial roles in intracellular degradation of sphingolipids [78]. Previously, we showed that PSAP levels in GBM-EVs have a significant, positive correlation to *in vitro* GBM cell invasion [25] and multiple studies have reported high PSAP expression in clinical glioma specimens, glioma-stem cells and cell-lines [79, 80] with elevated serum levels in advanced-stage prostate cancer patients [81]. While PSAP was not confidently identified in EVs purified from glioma (II-IV) neurosurgical fluids [24, 25], here PSAP levels are significantly lower in GBM plasma-EVs relative to controls. Other studies have proposed contrary roles for PSAP that include a tumour-secreted inhibitor of metastasis. In prostate cancer, high PSAP levels are associated with weakly- or non-metastatic tumours, creating microenvironments in distal tissues that are refractory to metastasis [82]. Although gliomas rarely metastasise beyond the CNS [83], the significant reduction of PSAP secretion in GBM/GIV plasma-EVs detected in this study may reflect a mechanism whereby PSAP levels are selectively retained in GBM tissue, to promote growth and invasion of GBM that is limited to the brain.

Interestingly, TOMM40 was measured in significantly high levels in plasma-EVs from GBM/GIV patient groups relative to HC (adj. *p-val* ≤0.05; Figure 6A,Supplementary Table 4 and 5). TOMM40 was also more highly expressed in GIV relative to GIII (adj. *p-val*≤0.05) and GII (borderline significance of adj. *p-val*=0.06; Supplementary File D). We previously reported significantly higher TOMM40 expression in EVs from GBM neurosurgical aspirates compared to GII-III glioma [24]. TOMM40 plays a central role in cell metabolism, shuttling proteins and acting as the gateway for protein entrance into the mitochondria [84]. High TOMM40 levels are associated with a super-invasive subtype of melanoma [84], with BRCA1/2 expression and mutations [85] and are a histological feature for the squamous subtype of non-small cell lung cancer [86]. Increased levels of TOMM40 in plasma-EV GBM/GIV samples relative to GII-GIII gliomas, as well as non-glioma and healthy controls, may be a useful distinguishing biomarker for GBM and should be further explored.

### T-complex protein 1 ring complex (TRiC) interactome may indicate GBM presence and/or progression

T-complex protein 1 ring complex (TRiC; also known as chaperonin containing TCP1 complex, CCT) protein subunits, CCT2, CCT3, CCT4, CCT5, CCT7 and TCP1 were measured in high levels (FC≥2) in plasma-EVs from GBM patients relative to the controls, with significant increases detected for CCT2 and CCT7 (adj. *p-val* ≤ 0.05, Supplementary File C and D). We previously reported significantly higher CCT2 and CCT7 levels in GBM neurosurgical aspirate-EVs compared to GII-III glioma, with analogous transcript expression levels and DNA copy numbers reported for CCT2 and CCT7 in GBM tissue compared to normal brain in TCGA datasets [24]. While the changes observed here are relative to healthy and non-glioma controls, the significantly high expression of TRiC components CCT2 and CCT7 within plasma-EVs of GBM/GIV patients, not observed for less aggressive glioma subtypes, might therefore relate to primary GBM biology (*IDH*-wt) and disease severity.

TRiC is a torus-shaped complex composed of eight distinct non-identical protein subunits (TCP1, CCT2, CCT3, CCT4, CCT5, CCT6A, CCT7, CCT8) [87, 88]. It is a cytoplasmic molecular chaperone that assists newly formed proteins to fold correctly [87, 89]. Increases in TRiC expression has been associated with tumorigenesis, influencing molecular pathways that contribute to tumour progression, such as p53 and STAT3 [87, 90-92]. Defects in TRiC activity are associated with protein misfolding and cytotoxicity [93] and have been observed in multiple neuropathologies [94-96]. The role of TRiC in tumour aggression has been further corroborated by the successful implementation of cytotoxic peptides such as CT20p that reduce tumour burden by targeting CCT2 [97]. CT20p inhibition of CCT2 reduces TRiC activity and alters the level of other TRiC components, interfering with the proper folding of oncogenic molecules such as STAT3 [97]. While TRiC is a highly functional chaperone as a complex, the subunits also have individual functions [98]. CCT7 expression is associated with cancer cell growth and maintenance [99], down-regulated during the DNA damage response in GBM [99] and is associated with poor clinical outcome in GBM patients [100]. CCT2 also has important functional links to cancer, including survival of breast cancer cells and negative survival associations in breast cancer [101], non-small cell lung cancer [102] and diffuse intrinsic pontine glioma [103].

TRiC’s folding activity requires assistance from other protein molecules that together comprise the TRiC interactome [89]. TRiC interactome proteins, PARK7 and RECQL, were also DE in plasma-EVs in this study. PARK7 was significantly increased in both GBM and GIV sample groups, relative to controls (Figure 5C-1, C-2, Supplementary Tables 4 and 5) and RECQL was one of the top-50 DE plasma-EV proteins, with high levels distinguishing GIV from GIII gliomas (10.7-fold, adj. *p-val* = 0.0157, Supplementary File D) as well as GBM/GIV from controls. PARK7 has important roles in protecting cells from stress and apoptosis by acting as an oxidative stress sensor and redox-sensitive chaperone [104]. It has been increasingly implicated with neurodegenerative diseases [105] and across multiple cancers, stimulating cell proliferation, cell invasion and cancer metastasis [106-108]. PARK7 promotes tumorigenesis by influencing anti-apoptotic and pro-survival pathways [106], and epithelial-mesenchymal transition [109, 110]. It is proposed to promote cancer progression by activating Akt/mTOR, MEK/ERK, NF-κB and HIFα signalling pathways, or repressing p53, JNK and ASK1 signalling pathways [111]. High PARK7 expression is associated with reactive astrocytes [106], confers poor prognosis in breast cancer [112], is found in more invasive breast cancer cells [110], is highly expressed in advanced-stage ovarian cancer [113] and has a positive correlation to recurrence and metastasis of lung cancer [114, 115]. PARK7 has been found in high concentrations in body fluids of cancer patients suggesting its potential as a non-invasive cancer biomarker [111]. In GBM, high cytoplasmic PARK7 levels are associated with strong nuclear p53 expression and inversely correlated with EGFR expression [106].

The RecQ helicase family is a group of highly conserved DNA unwinding enzymes involved in maintaining chromosomal instability [116]. RECQL-deficient cells have been shown to accumulate DNA damage, display sensitivity to DNA damaging agents and exhibit chromosomal instability [116, 117]. RECQL is amplified and overexpressed in many clinical cancer specimens and previously implicated in CNS tumours [116]. There are reports of RECQL overexpression in GBM [118], ovarian cancer [119] and head and neck squamous cell carcinoma [120]. It is associated with EMT [116], cell migration, cancer invasion and metastasis, as well as exhibiting predictive and prognostic biomarker potential [116]. High RECQL expression correlates to higher histological grade and ki-67 expression in hepatocellular carcinoma [121], high proliferation in ovarian cancer [119] and faster tumour progression and reduced survival in adenocarcinoma patients [122, 123]. In GBM, high expression of RECQ1 promotes cell proliferation, S-phase cell-cycle progression and γ-H2AX foci formation in GBM cell-lines [118]. Although these studies imply an association of RECQL with tumour progression and invasion, the molecular mechanisms by which RECQL supports this remains unknown. In addition, RECQL has also been identified in cell-derived memetic nanovesicles and exosomes derived from neuroblastoma [124].

Taken together, results here and in previous GBM-EV studies [24, 25], implicate a role for TRiC and its interactome in GBM pathology, which are reflected in EVs released into the circulation. It is possible that EV-associated TRiC and its interacting proteins could serve as robust glioma biomarkers with potential to indicate the presence of glioma and progression. Further investigation of TRiC components and its interactome in plasma-EVs is required. Building larger diffuse glioma cohorts will improve the power of future studies and allow for machine learning methods to train, validate and test potential glioma plasma-EV biomarkers. This will open avenues for building predictive models that stratify glioma patients and determine glioma progression and recurrence. Furthermore, serial blood sampling from glioma patients over the course of their disease and correlating plasma-EV profiles with clinical, pathological and radiological parameters will allow for predictive models to be designed based on a panel of select GBM/glioma biomarkers.

## Concluding remarks

With current tumour surveillance approaches, diffuse glioma progression can be confused with common treatment-related brain changes. This is particularly problematic as patients may continue to receive an ineffective treatment or have an effectual treatment stopped prematurely. The development of a liquid biopsy that is capable of monitoring glioma patients accurately and reliably over the course of their disease would significantly improve the clinical management of glioma. An ideal glioma liquid biopsy would diagnose and stratify glioma patients, and would be capable of monitoring glioma progression and treatment resistance, whilst distinguishing progression from common treatment-related effects. Using SWATH-MS, we have achieved the most comprehensive and in-depth proteomic coverage of plasma-EVs to-date and established a clear advantage of applying this analytical platform for blood-based EV biomarker discovery. Significant plasma-EV proteins were identified between glioma genetic-histological subtypes and grades, including proteins previously reported as markers of GBM invasion and aggression. Plasma-EV proteins reflect tumour progression and have the potential to describe diagnostic, prognostic and predictive biomarker signatures. Future studies analysing plasma-EV proteins in larger longitudinal cohorts of glioma patients may substantiate the markers determined in this study for the conceptualisation of a glioma liquid biopsy. Elucidating clinically-relevant EV biomarkers that offer non-invasive, early indications of tumour recurrence are likely to have significant clinical utility, opening avenues to evaluate glioma progression and treatment response and in turn, improve the clinical management of patients.

## Author contributions

All authors contributed to manuscript preparation and approved the submission of the work presented here. Specific contributions are as follows: S.H. performed the technical and analytical work including plasma processing, EV isolation, proteome preparation, nanosight, proteomics, *in silico* analyses, bioinformatics, data interpretation, and presentation. A.A. provided SWATH-MS experimental design, method development and spectral library data. H.W. and M.L. characterised clinical cases and provided technical expertise. N.H. developed R scripts for data analysis and data presentation. J.S. performed neuropathological assessments. B.S. and H.S. assisted with clinical sample procurement and case characterisation. M.E.B contributed to experimental design, neuropathological assessments and case characterisation. K.L.K provided overall experimental design, cohort characterisations, data interpretation, and presentation.

## Acknowldegments

Many thanks to the Royal Prince Alfred Hospital (RPAH) clinical staff, with special thanks to Mary Lordan and Jane Raftesath. Thank you to the Neuropathology Tumour and Tissue Bank at Royal Prince Alfred Hospital and Brain and Mind Centre for the provision of blood specimens and associated information. This work was enabled by access to several facilities of the University of Sydney, including the Bosch Institute Molecular Biology Facility, Sydney Mass Spectrometry, and Advanced Microscopy Facility. Thank you to Saeideh Ebrahimkhani for the transmission electron microscopy images.

