## Supplementary material for "A comprehensive proteomic SWATH-MS workflow for profiling blood extracellular vesicles: a new avenue for glioma tumour surveillance": Supplementary Material.pdf

**Table 1** - **Patient cohorts for SWATH-MS analysis of plasma-EVs.** Patients were categorised by genetic-histological tumour subtype, including *IDH-wt* glioblastoma/astrocytoma (GBM), *IDH-mut* astrocytoma (AST) and *IDH-mut* 1p19q codel oligodendroglioma (OLI), or by grade (G) II-IV. Control cohort included non-glioma, grade I meningioma (MEN) and healthy controls (HC). Plasma derived from patients with recurrent tumours are highlighted.

| Patient sample | Gender | Age | Diagnosis | IDH | 1p/19q | Matching patient recurrence | Sample category | |
|---|---|---|---|---|---|---|---|---|
| | | | | | | | Genetic-histologic | Grade |
| **GBM1** | M | 67 | GBM, WHO (2016), Grade IV | *wt* | - | | GBM | GIV |
| **GBM2** | M | 45 | GBM, WHO (2016), Grade IV | *wt* | - | | GBM | GIV |
| **GBM3** | F | 75 | GBM, WHO (2016), Grade IV | *wt* | - | | GBM | GIV |
| **GBM4** | M | 47 | GBM, WHO (2016), Grade IV | *wt* | - | | GBM | GIV |
| **GBM5** | F | 75 | GBM, WHO (2016), Grade IV | *wt* | - | | GBM | GIV |
| **GBM6** | M | 73 | GBM, WHO (2016), Grade IV | *wt* | - | | GBM | GIV |
| **GBM7** | M | 74 | GBM, WHO (2016), Grade IV | *wt* | - | | GBM | GIV |
| **GBM8** | M | 75 | GBM, WHO (2016), Grade IV | *wt* | - | | GBM | GIV |
| **GBM9** | F | 60 | GBM, WHO (2016), Grade IV | *wt* | - | | GBM | GIV |
| **GBM10** | F | 69 | GBM, WHO (2016), Grade IV | *wt* | - | | GBM | GIV |
| **GBM11** | F | 46 | GBM, WHO (2016), Grade IV | *wt* | - | | GBM | GIV |
| **GBM12** | F | 64 | GBM, WHO (2016), Grade IV | *wt* | - | | GBM | GIV |
| **GBM13** | M | 63 | GBM, WHO (2016), Grade IV | *wt* | - | | GBM | GIV |
| **GBM14** | M | 34 | GBM, WHO (2016), Grade IV | *wt* | - | | GBM | GIV |
| **GBM15** | M | 60 | GBM, WHO (2016), Grade IV | *wt* | - | | GBM | GIV |
| **GBM16** | M | 61 | GBM, WHO (2016), Grade IV | *wt* | - | | GBM | GIV |
| **GBM17** | F | 55 | GBM, WHO (2016), Grade IV | *wt* | - | | GBM | GIV |
| **GBM18** | M | 45 | GBM, WHO (2016), Grade IV | *wt* | - | | GBM | GIV |
| **GBM18.1** | M | 46 | GBM, WHO (2016), Grade IV | *wt* | - | GBM18 recurrence | GBM | GIV |
| **GBM19** | M | 33 | GBM, WHO (2016), Grade IV | *mut* | - | | AST | GIV |
| **GBM20 (AST10.1)** | M | 52 | GBM, WHO (2016), Grade IV | *mut* | - | AST11 progression | AST | GIV |
| **GBM21** | M | 72 | GBM, WHO (2016), Grade IV | *wt* | - | | GBM | GIV |
| **GBM22** | M | 72 | GBM, WHO (2016), Grade IV | *wt* | - | | GBM | GIV |
| **GBM23** | M | 33 | GBM, WHO (2016), Grade IV | *wt* | - | | GBM | GIV |
| **GBM24** | M | 63 | GBM, WHO (2016), Grade IV | *wt* | - | | GBM | GIV |
| **GBM25** | M | 85 | GBM, WHO (2016), Grade IV | *wt* | - | | GBM | GIV |
| **AST1** | F | 57 | Diffuse astrocytoma, WHO (2016), Grade II | *mut* | - | | AST | GII |
| **AST2** | M | 21 | Diffuse astrocytoma, WHO (2016), Grade II | *mut* | - | | AST | GII |
| **AST3** | F | 37 | Diffuse astrocytoma, WHO (2016), Grade II | *mut* | - | | AST | GII |
| **AST4** | M | 33 | Diffuse astrocytoma, WHO (2016), Grade II | *mut* | - | | AST | GII |
| **AST5** | M | 42 | Diffuse astrocytoma, WHO (2016), Grade II | *mut* | - | | AST | GII |
| **AST5.1** | M | 44 | Anaplastic astrocytoma, WHO (2016), Grade III | *mut* | - | AST5 progression | AST | GIII |
| **AST6** | M | 68 | Anaplastic astrocytoma, WHO (2016), Grade III | *mut* | - | | AST | GIII |
| **AST7** | F | 27 | Anaplastic astrocytoma, WHO (2016), Grade III | *mut* | - | | AST | GIII |
| **AST8** | M | 25 | Anaplastic astrocytoma, WHO (2016), Grade III | *mut* | - | | AST | GIII |
| **AST9** | M | 31 | Anaplastic astrocytoma, WHO (2016), Grade III | *mut* | - | | AST | GIII |
| **AST10** | M | 51 | Anaplastic astrocytoma, WHO (2016), Grade III | *mut* | - | | AST | GIII |
| **OLI1** | M | 34 | Oligodendroglioma, WHO (2016), Grade II | *mut* | codel | | OLI | GII |
| **OLI2** | M | 33 | Anaplastic oligodendroglioma, WHO (2016), Grade III | *mut* | codel | | OLI | GIII |
| **OLI3** | M | 38 | Anaplastic oligodendroglioma, WHO (2016), Grade III | *mut* | codel | | OLI | GIII |
| **OLI4** | M | 41 | Anaplastic oligodendroglioma, WHO (2016), Grade III | *mut* | codel | | OLI | GIII |
| **MEN1** | F | 65 | Meningothelial meningioma, WHO (2016), Grade I | - | - | | MEN | |
| **MEN2** | F | 48 | Meningothelial meningioma, WHO (2016), Grade I | - | - | | MEN | |
| **MEN3** | M | 59 | Meningothelial meningioma, WHO (2016), Grade I | - | - | | MEN | |
| **MEN4** | M | 45 | Meningothelial meningioma, WHO (2016), Grade I | - | - | | MEN | |
| **MEN5** | M | 71 | Meningioma, WHO (2016), Grade I | - | - | | MEN | |
| **HC1** | M | 50 | - | - | - | | HC | |
| **HC2** | F | 55 | - | - | - | | HC | |
| **HC3** | F | 47 | - | - | - | | HC | |
| **HC4** | M | 38 | - | - | - | | HC | |
| **HC5** | M | 38 | - | - | - | | HC | |
| **HC6** | F | 53 | - | - | - | | HC | |

**Table 2-** Pooled peptides used for the generation of a comprehensive custom spectral library. Peptides from primary GBM cells, neurosurgical aspirate EVs, GBM tumour microsomal and soluble samples, K562 cells and fresh frozen paraffin embedded (FFPE) skin cancers were fractionated by hydrophilic interaction liquid chromatography (HILIC) on an Agilent HPLC system.

| **Sample type** | **Peptide samples** | **Pooled peptides for HILIC fractionation (µg)** | **Number of HILIC fractions** |
|---|---|---|---|
| ***Primary GBM cells*** | JK2 cell peptides<br>WK1 cell peptides | 20 µg | 8 |
| ***Enriched-neurosurgical aspirate EVs*** | Glioblastoma, grade IV glioma (*n*=19)<br>Grade II-III glioma (*n*=8)<br>Benign, Grade I Meningioma (*n*=5) | 20 µg | 10 |
| ***GBM tumour microsomal and soluble samples*** | Peptide pool of six specimens (3 tumour pairs; primary and recurrent GBM) of microsomal and soluble proteomes | 20 µg | 10 |
| ***K562 cell*** | K562 cell peptides | 20 µg | 8 |
| ***FFPE skin*** | Normal skin (*n*=12)<br>Actinic keratosis (*n*=18)<br>Bowen's disease (*n*=19)<br>Cutaneous squamous-cell carcinoma (cSCC; *n*=44) | 20 µg | 6 |

**Table 3 –** The 159 variable sequential precursor isolation windows defined by the SWATH Variable Window Calculator, with a *m/z* range of 350-1750 and a minimum window width of 3 Da.

| SWATH Window | Start mass (Da) | Stop mass (Da) | CES | SWATH Window | Start mass (Da) | Stop mass (Da) | CES | SWATH Window | Start mass (Da) | Stop mass (Da) | CES |
|---|---|---|---|---|---|---|---|---|---|---|---|
| **1** | 349.5 | 352.5 | 5 | **54** | 455.5 | 458.5 | 5 | **107** | 561.6 | 566 | 5 |
| **2** | 351.5 | 354.5 | 5 | **55** | 457.5 | 460.5 | 5 | **108** | 565 | 569.4 | 5 |
| **3** | 353.5 | 356.5 | 5 | **56** | 459.5 | 462.5 | 5 | **109** | 568.4 | 573.7 | 5 |
| **4** | 355.5 | 358.5 | 5 | **57** | 461.5 | 464.5 | 5 | **110** | 572.7 | 577.1 | 5 |
| **5** | 357.5 | 360.5 | 5 | **58** | 463.5 | 466.5 | 5 | **111** | 576.1 | 581.3 | 5 |
| **6** | 359.5 | 362.5 | 5 | **59** | 465.5 | 468.5 | 5 | **112** | 580.3 | 585.6 | 5 |
| **7** | 361.5 | 364.5 | 5 | **60** | 467.5 | 470.5 | 5 | **113** | 584.6 | 589.8 | 5 |
| **8** | 363.5 | 366.5 | 5 | **61** | 469.5 | 472.5 | 5 | **114** | 588.8 | 594.1 | 5 |
| **9** | 365.5 | 368.5 | 5 | **62** | 471.5 | 474.5 | 5 | **115** | 593.1 | 599.2 | 5 |
| **10** | 367.5 | 370.5 | 5 | **63** | 473.5 | 476.5 | 5 | **116** | 598.2 | 603.4 | 5 |
| **11** | 369.5 | 372.5 | 5 | **64** | 475.5 | 478.5 | 5 | **117** | 602.4 | 608.5 | 5 |
| **12** | 371.5 | 374.5 | 5 | **65** | 477.5 | 480.5 | 5 | **118** | 607.5 | 612.7 | 5 |
| **13** | 373.5 | 376.5 | 5 | **66** | 479.5 | 482.5 | 5 | **119** | 611.7 | 617.8 | 5 |
| **14** | 375.5 | 378.5 | 5 | **67** | 481.5 | 484.5 | 5 | **120** | 616.8 | 622.9 | 5 |
| **15** | 377.5 | 380.5 | 5 | **68** | 483.5 | 486.5 | 5 | **121** | 621.9 | 628 | 5 |
| **16** | 379.5 | 382.5 | 5 | **69** | 485.5 | 488.5 | 5 | **122** | 627 | 633.1 | 5 |
| **17** | 381.5 | 384.5 | 5 | **70** | 487.5 | 490.5 | 5 | **123** | 632.1 | 639.1 | 5 |
| **18** | 383.5 | 386.5 | 5 | **71** | 488.6 | 491.6 | 5 | **124** | 638.1 | 645 | 5 |
| **19** | 385.5 | 388.5 | 5 | **72** | 490.3 | 493.3 | 5 | **125** | 644 | 651 | 5 |
| **20** | 387.5 | 390.5 | 5 | **73** | 492 | 495 | 5 | **126** | 650 | 656.1 | 5 |
| **21** | 389.5 | 392.5 | 5 | **74** | 493.7 | 496.7 | 5 | **127** | 655.1 | 661.2 | 5 |
| **22** | 391.5 | 394.5 | 5 | **75** | 495.4 | 498.4 | 5 | **128** | 660.2 | 665.4 | 5 |
| **23** | 393.5 | 396.5 | 5 | **76** | 497.1 | 500.1 | 5 | **129** | 664.4 | 670.5 | 5 |
| **24** | 395.5 | 398.5 | 5 | **77** | 498.8 | 501.8 | 5 | **130** | 669.5 | 674.8 | 5 |
| **25** | 397.5 | 400.5 | 5 | **78** | 500.5 | 503.5 | 5 | **131** | 673.8 | 679.9 | 5 |
| **26** | 399.5 | 402.5 | 5 | **79** | 502.2 | 505.2 | 5 | **132** | 678.9 | 685 | 5 |
| **27** | 401.5 | 404.5 | 5 | **80** | 503.9 | 506.9 | 5 | **133** | 684 | 690.1 | 5 |
| **28** | 403.5 | 406.5 | 5 | **81** | 505.6 | 508.6 | 5 | **134** | 689.1 | 695.2 | 5 |
| **29** | 405.5 | 408.5 | 5 | **82** | 507.3 | 510.3 | 5 | **135** | 694.2 | 700.2 | 5 |
| **30** | 407.5 | 410.5 | 5 | **83** | 509 | 512 | 5 | **136** | 699.2 | 705.3 | 5 |
| **31** | 409.5 | 412.5 | 5 | **84** | 510.7 | 513.7 | 5 | **137** | 704.3 | 710.4 | 5 |
| **32** | 411.5 | 414.5 | 5 | **85** | 512.4 | 515.4 | 5 | **138** | 709.4 | 716.4 | 5 |
| **33** | 413.5 | 416.5 | 5 | **86** | 514.1 | 517.1 | 5 | **139** | 715.4 | 722.3 | 5 |
| **34** | 415.5 | 418.5 | 5 | **87** | 515.8 | 518.8 | 5 | **140** | 721.3 | 728.3 | 5 |
| **35** | 417.5 | 420.5 | 5 | **88** | 517.5 | 520.5 | 5 | **141** | 727.3 | 734.2 | 5 |
| **36** | 419.5 | 422.5 | 5 | **89** | 519.2 | 522.2 | 5 | **142** | 733.2 | 740.2 | 5 |
| **37** | 421.5 | 424.5 | 5 | **90** | 520.9 | 523.9 | 5 | **143** | 739.2 | 746.1 | 5 |
| **38** | 423.5 | 426.5 | 5 | **91** | 522.6 | 525.6 | 5 | **144** | 745.1 | 752.9 | 5 |
| **39** | 425.5 | 428.5 | 5 | **92** | 524.2 | 527.2 | 5 | **145** | 751.9 | 758.9 | 5 |
| **40** | 427.5 | 430.5 | 5 | **93** | 525.9 | 528.9 | 5 | **146** | 757.9 | 765.7 | 5 |
| **41** | 429.5 | 432.5 | 5 | **94** | 527.6 | 530.6 | 5 | **147** | 764.7 | 771.6 | 5 |
| **42** | 431.5 | 434.5 | 5 | **95** | 529.3 | 532.3 | 5 | **148** | 770.6 | 778.4 | 5 |
| **43** | 433.5 | 436.5 | 5 | **96** | 531 | 534.6 | 5 | **149** | 777.4 | 788.6 | 5 |
| **44** | 435.5 | 438.5 | 5 | **97** | 533.6 | 537.1 | 5 | **150** | 787.6 | 802.2 | 5 |
| **45** | 437.5 | 440.5 | 5 | **98** | 536.1 | 539.1 | 5 | **151** | 801.2 | 821.7 | 5 |
| **46** | 439.5 | 442.5 | 5 | **99** | 537.8 | 541.4 | 5 | **152** | 820.7 | 845.5 | 5 |
| **47** | 441.5 | 444.5 | 5 | **100** | 540.4 | 544.8 | 5 | **153** | 844.5 | 871 | 5 |
| **48** | 443.5 | 446.5 | 5 | **101** | 543.8 | 547.3 | 5 | **154** | 870 | 907.5 | 5 |
| **49** | 445.5 | 448.5 | 5 | **102** | 546.3 | 550.7 | 5 | **155** | 906.5 | 965.3 | 5 |
| **50** | 447.5 | 450.5 | 5 | **103** | 549.7 | 553.3 | 5 | **156** | 964.3 | 1021.4 | 5 |
| **51** | 449.5 | 452.5 | 5 | **104** | 552.3 | 556.7 | 5 | **157** | 1020.4 | 1209.1 | 5 |
| **52** | 451.5 | 454.5 | 5 | **105** | 555.7 | 560.1 | 5 | **158** | 1208.1 | 1749.4 | 5 |
| **53** | 453.5 | 456.5 | 5 | **106** | 559.1 | 562.6 | 5 | **159** | 1748.4 | 1751.4 | 5 |

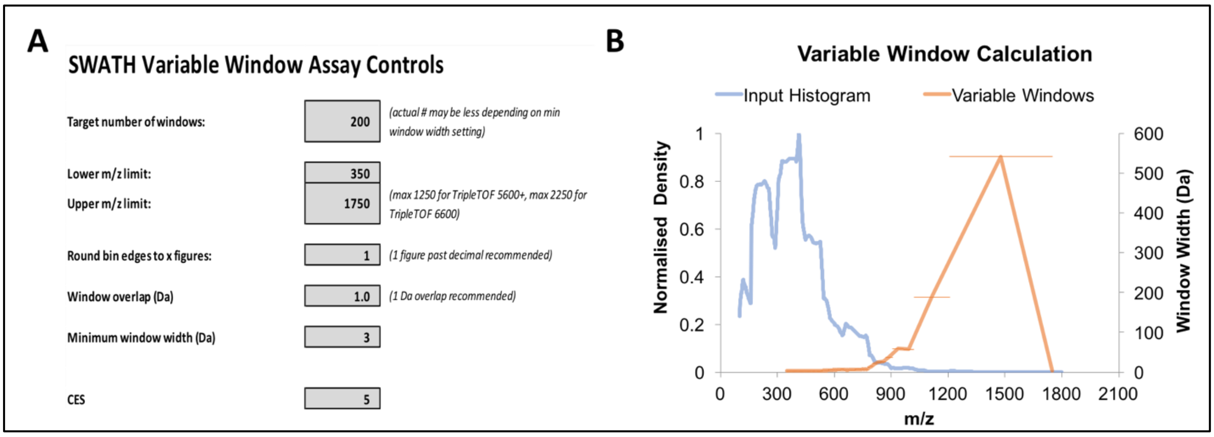

**Figure 1 -** (A) Parameters used to generate SWATH variable windows. (B) Graph of the *m/z* histogram and the variable swath windows that were generated. Smaller *m/z* window ranges were generated for higher inputs.

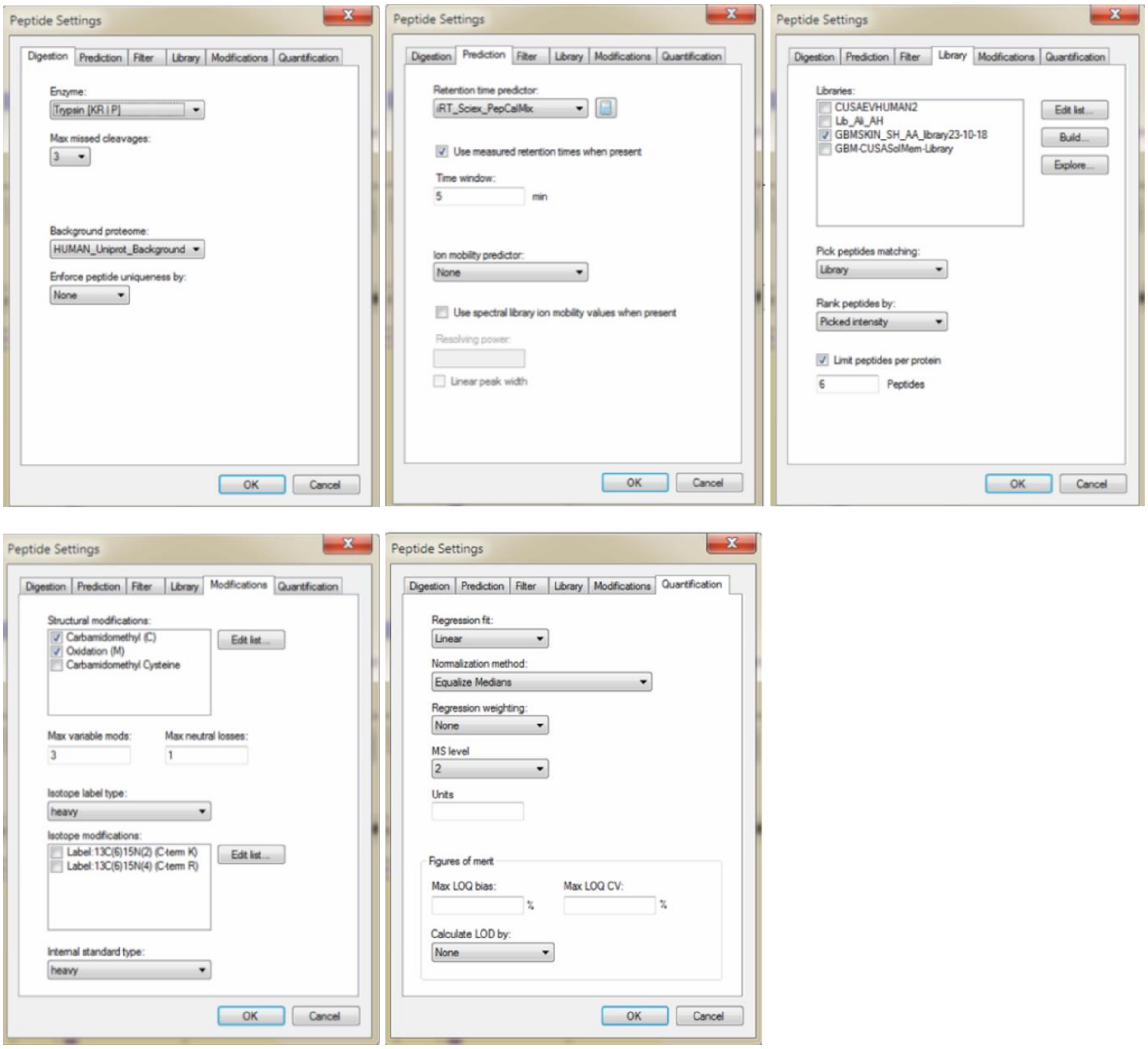

**Figure 2 -** Peptide settings used for the analysis of SWATH-MS data in Skyline4.2.

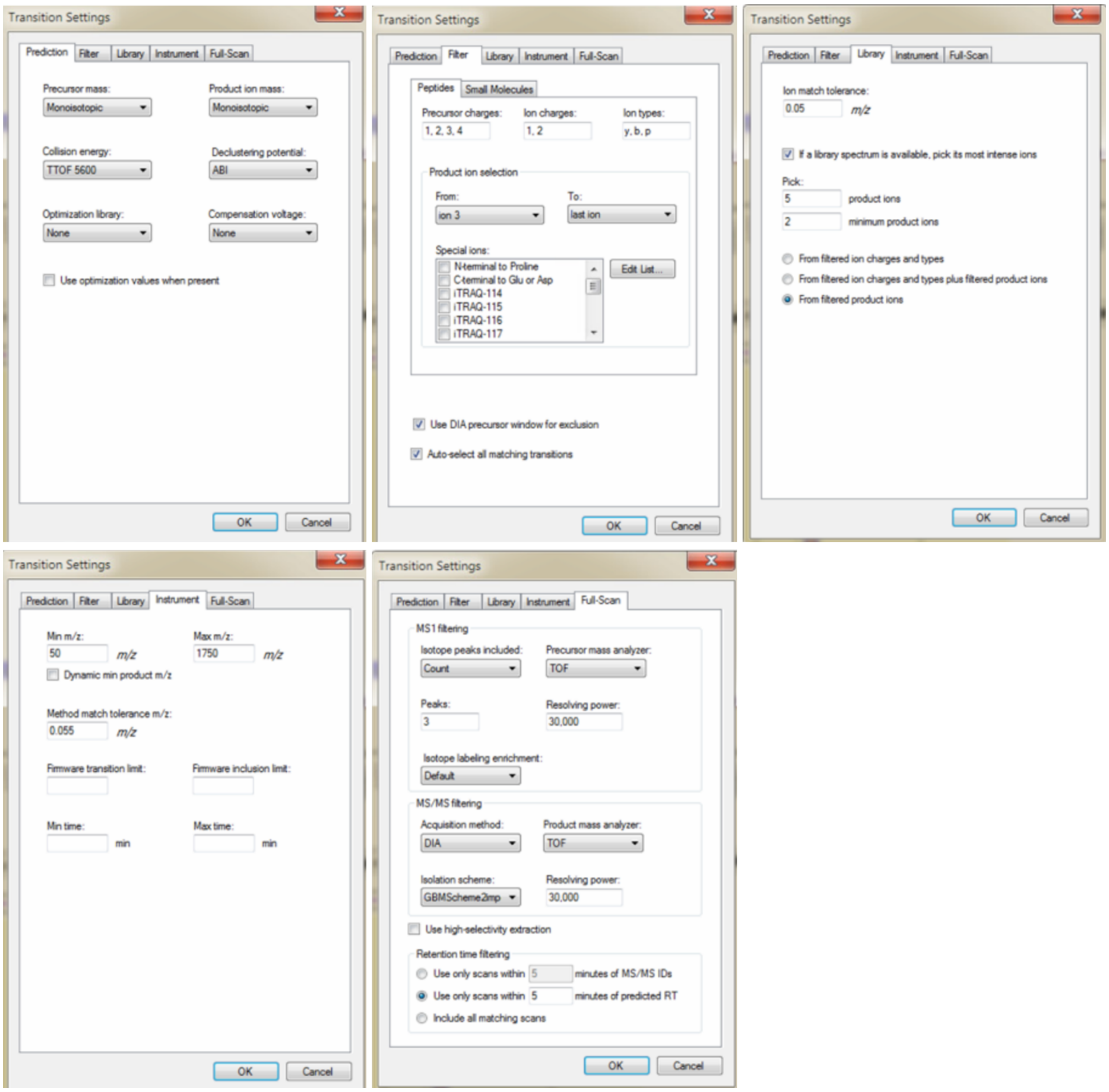

**Figure 3 -** Transition settings used for the analysis of SWATH data in Skyline4.2.

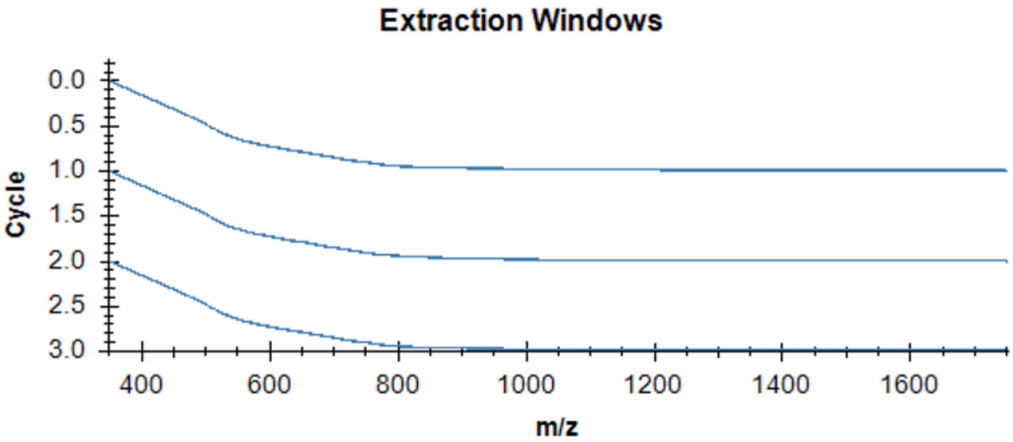

**Figure 4 -** Skyline4.2 isolation scheme of the variable isolation windows used on the TripleTOF®6600 to acquire SWATH-MS data.

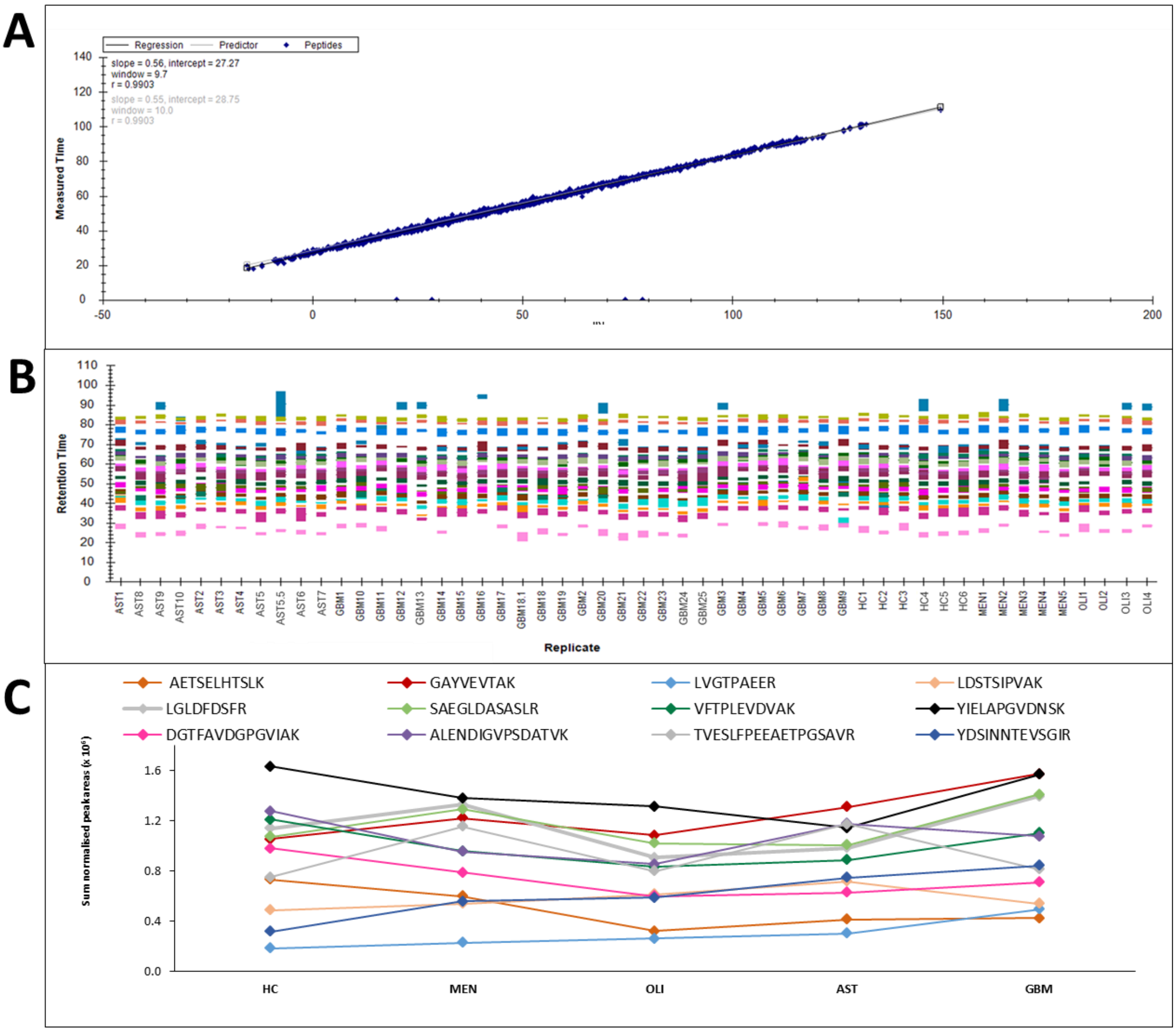

**Figure 5 - Data quality and reproducibility for glioma plasma-EVs**. (A) iRT calculator for the 20 *PepCalMix* peptides to allow retention time prediction and alignment of the SWATH-MS files to the spectral library. The *PepCalMix* peptides spiked-in to the spectral library were calibrated to the *PepCalMix* iRT definition values in Skyline4.2. The iRT linear regression between the measured retention times of the peptides (y-axis) versus their respective iRT definition values (x-axis) had a Pearson's correlation coefficient (*r*) of 0.9903. (B) The calibrated retention time ranges for 20 *PepCalMix* peptides for plasma EV samples analysed by SWATH-MS. (C) Scatter plot of the total peak area assigned to 12 *PepCalMix* peptides for patient plasma EVs categorised by genetic-histological subtype.

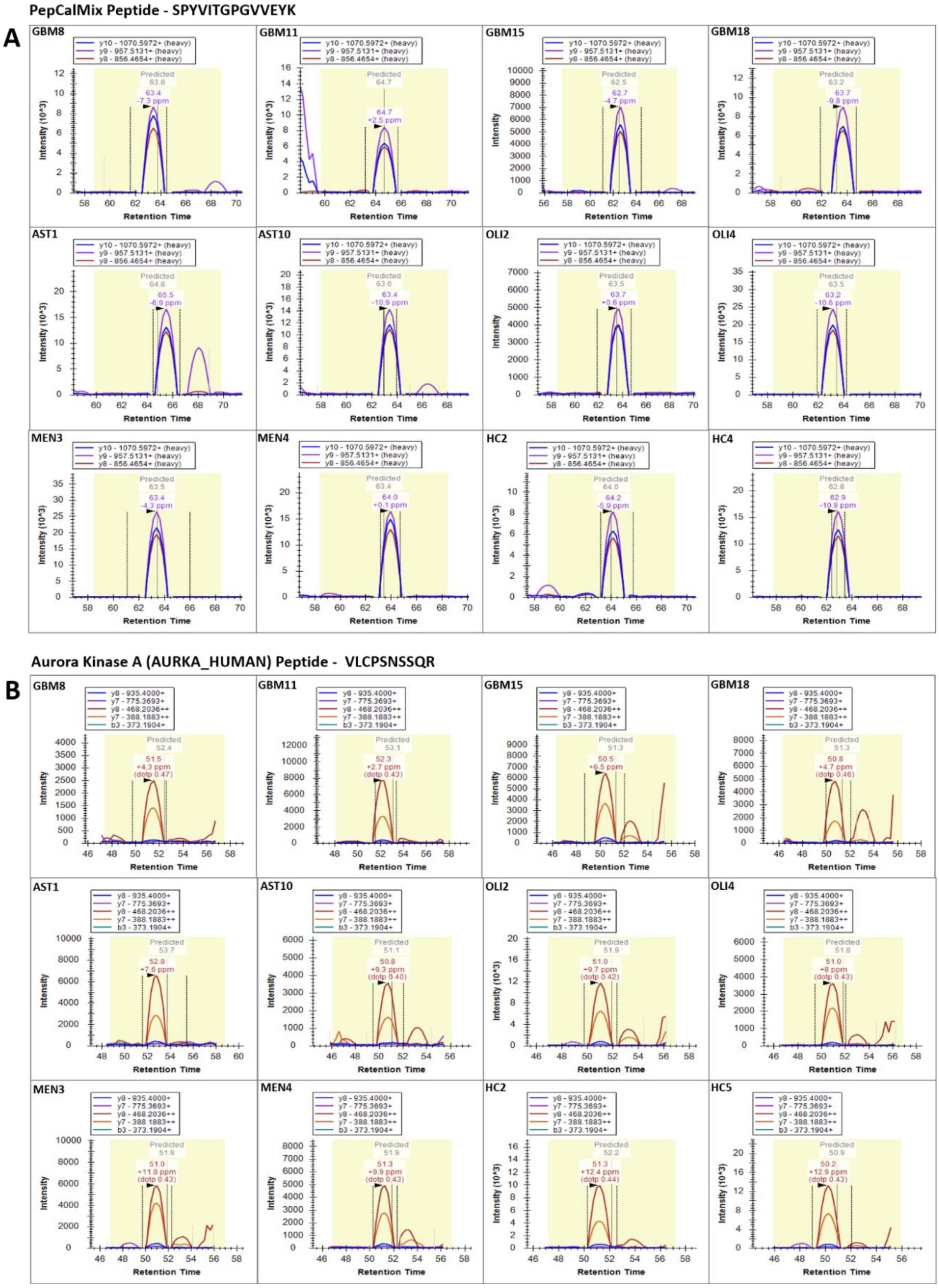

**Figure 6 - Extracted ion chromatograms (XIC) of standard *PepCalMix* and Aurora Kinase A peptide.** (A) XIC for the PepCalMix peptide SPYVITGPGVVEYK, and its product ions y8 (856.4654+; red), y9 (957.5131+; purple) and y10 (1070.5972+; red) across 12 plasma EV samples analysed by SWATH-MS. (B) The XIC for Aurora Kinase A (AURKA_HUMAN) peptide VLCPSNSSQR (574.2799 +++ (dotp 0.38)) and its product ions y8 (935.4000+; blue), y7 (7875.3693+; purple), y8 (468.2036++; red), y7 (388.1883++; orange) and b3 (373.1904+; green) across the same 12 samples.

**A**

| *Top canonical pathways* | *p-value* | *Overlap* |
|---|---|---|
| Synaptogenesis Signaling Pathway | $1.06 \times 10^{-8}$ | 8.5% (26/307) |
| Virus Entry via Endocytic Pathways | $3.32 \times 10^{-5}$ | 10.3% (11/107) |
| Acute Phase Response Signaling | $6.09 \times 10^{-5}$ | 7.9% (14/178) |
| Huntington's Disease Signaling | $9.97 \times 10^{-5}$ | 6.8% (16/234) |
| Clathrin-mediated Endocytosis Signaling | $1.37 \times 10^{-4}$ | 7.3% (14/192) |
| ***Diseases and disorders*** | ***p-value*** | ***#Molecules*** |
| Cancer | $2.28 \times 10^{-2} – 8.10 \times 10^{-19}$ | 450 |
| Endocrine System Disorders | $2.28 \times 10^{-2} – 8.10 \times 10^{-19}$ | 388 |
| Organismal Injury and Abnormalities | $2.28 \times 10^{-2} – 8.10 \times 10^{-19}$ | 455 |
| Inflammatory response | $2.28 \times 10^{-2} – 1.64 \times 10^{-11}$ | 78 |
| Gastrointestinal disease | $2.28 \times 10^{-2} – 2.77 \times 10^{-9}$ | 403 |
| ***Molecular and cellular functions*** | **p-value** | ***#Molecules*** |
| Cellular compromise | $2.28 \times 10^{-2} – 1.64 \times 10^{-11}$ | 48 |
| Cellular assembly and organisation | $2.28 \times 10^{-2} – 1.01 \times 10^{-10}$ | 75 |
| Cellular function and maintenance | $2.07 \times 10^{-2} – 8.67 \times 10^{-9}$ | 104 |
| Protein synthesis | $2.05 \times 10^{-2} – 6.42 \times 10^{-5}$ | 80 |
| Post-translational modification | $2.28 \times 10^{-2} – 3.19 \times 10^{-4}$ | 24 |

**B**

| **Biological Process** | **Over/under representation** | **Fold Enrichment** | **p-val (FDR<0.05)** |
|---|---|---|---|
| ATP synthesis coupled proton transport (GO:0015986) | + | 6.06 | 0.0491 |
| Carboxylic acid biosynthetic process (GO:0046394) | + | 4.99 | 0.0194 |
| Organelle localization (GO:0051640) | + | 4.72 | 0.0339 |
| Cellular amino acid metabolic process (GO:0006520) | + | 4.65 | 0.0246 |
| Protein folding (GO:0006457) | + | 3.88 | 0.0352 |
| Cytoskeleton organization (GO:0007010) | + | 3.42 | 0.00507 |
| Formation of translation initiation ternary complex (GO:0001677) | + | 3.35 | 0.00879 |
| Translational termination (GO:0006415) | + | 3.35 | 0.008 |
| Translational elongation (GO:0006414) | + | 3.35 | 0.00733 |
| Vesicle-mediated transport (GO:0016192) | + | 2.12 | 0.0222 |
| Regulation of transcription by RNA polymerase II (GO:0006357) | - | 0.21 | 0.0496 |
| Transcription by RNA polymerase II (GO:0006366) | - | 0.18 | 0.00232 |
| **Reactome pathways** | **Over/under representation** | **Fold Enrichment** | **p-val (FDR<0.05)** |
| Retrograde neurotrophin signalling (R-HSA-177504) | + | 12.98 | 4.00E-02 |
| WNT5A-dependent internalization of FZD4 (R-HSA-5099900) | + | 12.12 | 4.35E-02 |
| EPHB-mediated forward signaling (R-HSA-3928662) | + | 7.76 | 1.10E-02 |
| Cargo recognition for clathrin-mediated endocytosis (R-HSA-8856825) | + | 4.17 | 3.86E-02 |
| MHC class II antigen presentation (R-HSA-2132295) | + | 4 | 1.86E-02 |
| Response to elevated platelet cytosolic Ca2+ (R-HSA-76005) | + | 3.79 | 2.42E-02 |
| Fcgamma receptor (FCGR) dependent phagocytosis (R-HSA-2029480) | + | 3.71 | 1.93E-02 |
| Innate Immune System (R-HSA-168249) | + | 2.47 | 1.60E-07 |
| Neutrophil degranulation (R-HSA-6798695) | + | 3.61 | 2.74E-08 |
| Intra-Golgi and retrograde Golgi-to-ER traffic (R-HSA-6811442) | + | 3.6 | 3.99E-03 |
| ER to Golgi Anterograde Transport (R-HSA-199977) | + | 3.54 | 2.41E-02 |
| Translation (R-HSA-72766) | + | 2.79 | 1.77E-02 |
| RHO GTPase Effectors (R-HSA-195258) | + | 2.78 | 2.33E-02 |
| Infectious disease (R-HSA-5663205) | + | 2.46 | 3.84E-02 |
| Vesicle-mediated transport (R-HSA-5653656) | + | 3.03 | 2.94E-08 |

**Figure 7 –** (A) Functional pathway analysis of significantly changed proteins (fold change ≥ 2, adj. p-val ≤0.05) in GBM circulating EVs relative to healthy controls. Top canonical pathways, diseases and disorders and molecular and cellular function are listed with the numbers of overlapping molecules and significance of associations (right-tailed Fisher exact test, *p* value). (B) PANTHER over-representation test results for biological processes and reactome pathways. Significant results (fold change ≥ 2, p-val ≤0.05) are displayed for proteins in plasma-EVs that are over/under-represented relative to the number of proteins expected in the respective PANTHER category for plasma-EVs.

**Table 4 – Glioma-specific protein changes in plasma EVs associated to histologic-genetic glioma subtype.** DE proteins are relative to both healthy and non-glioma controls. Proteins that are restricted to glioma samples are denoted by '+INF' and proteins that are restricted to controls (HC or MEN) are denoted by '-INF'.

| DE Proteins in *IDH-wt* GBM | | | | | | | | | | | | | | |
|---|---|---|---|---|---|---|---|---|---|---|---|---|---|---|
| | | | **Fold Change Relative to HC** | | | | | | **Fold Change Relative to MEN** | | | | | |
| **Protein** | | | ***IDH-mut* OLI** | | ***IDH-mut* AST** | | ***IDH-wt* GBM** | | ***IDH-mut* OLI** | | ***IDH-mut* AST** | | ***IDH-wt* GBM** | |
| **Accession** | **Gene Name** | **Name** | **FC** | ***p*-val** | **FC** | ***p*-val** | **FC** | ***p*-val** | **FC** | ***p*-val** | **FC** | ***p*-val** | **FC** | ***p*-val** |
| 1A03_HUMAN | HLA-A | HLA class I histocompatibility antigen, A-3 alpha chain | -5.74 | 0.913 | -142.53 | 0.222 | -539.20 | 0.005 | -6.55 | 0.977 | -162.48 | 0.218 | -614.68 | 0.008 |
| SAP_HUMAN | PSAP | Prosaposin | -16.17 | 0.854 | -259.70 | 0.037 | -276.91 | 0.005 | -7.27 | 0.977 | -116.79 | 0.104 | -124.53 | 0.016 |
| HEM4_HUMAN | UROS | Uroporphyrinogen-III synthase | -5.47 | 0.913 | -16.42 | 0.579 | -232.81 | 0.009 | -34.79 | 0.977 | -104.39 | 0.218 | -1480.5 | 0.001 |
| CO6_HUMAN | C6 | Complement component C6 | -12.56 | 0.887 | -6.15 | 0.755 | -231.54 | 0.015 | -6.82 | 0.977 | -3.34 | 0.894 | -125.82 | 0.046 |
| RPN1_HUMAN | RPN1 | Dolichyl-diphosphooligosaccharide--protein glycosyltransferase subunit 1 | -23.80 | 0.854 | -96.20 | 0.049 | -174.51 | 0.005 | -7.30 | 0.977 | -29.51 | 0.270 | -53.53 | 0.039 |
| DQB1_HUMAN | HLA-DQB1 | HLA class II histocompatibility antigen, DQ beta 1 chain | -15.40 | 0.854 | -9.31 | 0.579 | -144.01 | 0.003 | -40.33 | 0.696 | -24.38 | 0.247 | -377.09 | 0.000 |
| CE170_HUMAN | CEP170 | Centrosomal protein of 170 kDa | -19.69 | 0.328 | -85.68 | 0.016 | -107.19 | 0.001 | -3.68 | 0.894 | -16.02 | 0.109 | -20.04 | 0.046 |
| TOPK_HUMAN | PBK | Lymphokine-activated killer T-cell-originated protein kinase | -37.74 | 0.663 | -112.45 | 0.134 | -103.53 | 0.037 | +INF | 0.000 | +INF | 0.000 | +INF | 0.000 |
| ACYP2_HUMAN | ACYP2 | Acylphosphatase-2 | -4.63 | 0.913 | -15.46 | 0.579 | -86.40 | 0.033 | -13.41 | 0.977 | -44.78 | 0.270 | -250.32 | 0.009 |
| NUF2_HUMAN | NUF2 | Kinetochore protein Nuf2 | -4.25 | 0.403 | -3.02 | 0.272 | -48.67 | 0.000 | -2.91 | 0.825 | -2.07 | 0.533 | -33.35 | 0.002 |
| ENOG_HUMAN | ENO2 | Gamma-enolase | -1.35 | 0.988 | -6.88 | 0.633 | -48.49 | 0.033 | -2.13 | 0.996 | -10.85 | 0.583 | -76.46 | 0.024 |
| EPS8_HUMAN | EPS8 | Epidermal growth factor receptor kinase substrate 8 | -2.92 | 0.744 | -19.02 | 0.071 | -23.85 | 0.008 | -3.48 | 0.794 | -22.65 | 0.064 | -28.41 | 0.016 |
| STIM1_HUMAN | STIM1 | Stromal interaction molecule 1 | -2.57 | 0.826 | -5.39 | 0.267 | -15.88 | 0.007 | -1.87 | 0.984 | -3.91 | 0.439 | -11.51 | 0.033 |
| ITA7_HUMAN | ITGA7 | Integrin alpha-7 | -6.51 | 0.641 | -3.29 | 0.510 | -12.38 | 0.028 | -7.70 | 0.793 | -3.89 | 0.496 | -14.65 | 0.036 |
| ACSF2_HUMAN | ACSF2 | Acyl-CoA synthetase family member 2, mitochondrial | -4.56 | 0.663 | -3.45 | 0.458 | -11.77 | 0.009 | -7.17 | 0.617 | -5.42 | 0.233 | -18.48 | 0.003 |
| HIP1_HUMAN | HIP1 | Huntingtin-interacting protein 1 | -1.86 | 0.967 | -7.94 | 0.215 | -10.25 | 0.038 | -1.48 | 0.975 | -6.31 | 0.111 | -8.14 | 0.028 |
| PELO_HUMAN | PELO | Protein pelota homolog | -1.63 | 0.849 | -6.77 | 0.023 | -7.87 | 0.003 | -1.42 | 0.938 | -5.89 | 0.077 | -6.84 | 0.011 |
| SVIP_HUMAN | SVIP | Small VCP/p97-interacting protein | 1.135 | 0.980 | -1.788 | 0.605 | -7.10 | 0.035 | -1.005 | 0.995 | -2.041 | 0.545 | -8.10 | 0.022 |
| BD1L1_HUMAN | BOD1L1 | Biorientation of chromosomes in cell division protein 1-like 1 | 1.07 | 0.987 | -7.19 | 0.192 | -6.94 | 0.040 | -1.49 | 0.968 | -11.45 | 0.085 | -11.05 | 0.022 |
| NGLY1_HUMAN | NGLY1 | Peptide-N(4)-(N-acetyl-beta-glucosaminyl)asparagine amidase | 1.49 | 0.860 | -6.49 | 0.081 | -5.56 | 0.027 | -1.05 | 0.992 | -10.13 | 0.011 | -8.68 | 0.004 |
| HSP7E_HUMAN | HSPA14 | Heat shock 70 kDa protein 14 | -1.86 | 0.713 | -3.59 | 0.095 | -5.27 | 0.005 | -1.95 | 0.793 | -3.74 | 0.098 | -5.50 | 0.016 |
| WDR61_HUMAN | WDR61 | WD repeat-containing protein 61 | -2.45 | 0.511 | -2.87 | 0.133 | -4.34 | 0.005 | -2.65 | 0.825 | -3.10 | 0.086 | -4.70 | 0.008 |
| TRI14_HUMAN | TRIM14 | Tripartite motif-containing protein 14 | -2.94 | 0.545 | -2.00 | 0.510 | -4.21 | 0.028 | -3.51 | 0.622 | -2.38 | 0.439 | -5.02 | 0.032 |
| SART3_HUMAN | SART3 | Squamous cell carcinoma antigen recognized by T-cells 3 | 1.28 | 0.975 | -4.01 | 0.149 | -4.03 | 0.044 | -2.27 | 0.914 | -11.58 | 0.002 | -11.66 | 0.000 |
| DGKA_HUMAN | DGKA | Diacylglycerol kinase alpha | -1.54 | 0.900 | -2.00 | 0.402 | -3.34 | 0.041 | -1.72 | 0.928 | -2.23 | 0.367 | -3.73 | 0.047 |
| RM27_HUMAN | MRPL27 | 39S ribosomal protein L27, mitochondrial | 1.93 | 0.801 | -2.24 | 0.345 | -3.04 | 0.040 | 1.09 | 0.998 | -3.97 | 0.126 | -5.39 | 0.003 |
| LRCC7_HUMAN | LRRC7 | Leucine-rich repeat-containing protein 7 | -INF | 0.000 | 2.82 | 0.163 | 3.14 | 0.026 | NA | NA | +INF | 0.000 | +INF | 0.000 |
| ATIF1_HUMAN | ATP5IF1 | ATPase inhibitor, mitochondrial | -5.98 | 0.490 | 3.79 | 0.190 | 5.46 | 0.024 | -3.01 | 0.945 | 7.53 | 0.045 | 10.84 | 0.003 |
| APC1_HUMAN | ANAPC1 | Anaphase-promoting complex subunit 1 | -1.98 | 0.904 | 5.49 | 0.043 | 6.08 | 0.004 | -1.61 | 0.990 | 6.75 | 0.059 | 7.47 | 0.012 |
| ARPC3_HUMAN | ARPC3 | Actin-related protein 2/3 complex subunit 3 | 11.44 | 0.502 | 4.08 | 0.352 | 6.63 | 0.044 | 18.41 | 0.776 | 6.57 | 0.116 | 10.67 | 0.022 |
| TCPH_HUMAN | CCT7 | T-complex protein 1 subunit eta | -1.79 | 0.849 | 3.43 | 0.240 | 7.39 | 0.006 | -1.95 | 0.938 | 3.15 | 0.351 | 6.78 | 0.025 |
| CPTP_HUMAN | CPTP | Ceramide-1-phosphate transfer protein | 2.07 | 0.809 | 5.46 | 0.022 | 7.60 | 0.001 | 1.51 | 0.964 | 3.97 | 0.094 | 5.52 | 0.013 |
| WIPI3_HUMAN | WDR45B | WD repeat domain phosphoinositide-interacting protein 3 | -2.71 | 0.776 | 4.51 | 0.151 | 7.70 | 0.007 | -1.85 | 0.964 | 6.61 | 0.094 | 11.28 | 0.013 |

| | | | | | | | | | | | | | | |
|---|---|---|---|---|---|---|---|---|---|---|---|---|---|---|
| ANLN_HUMAN | ANLN | Anillin | -1.09 | 1.000 | 5.06 | 0.302 | 9.39 | 0.027 | 1.36 | 0.985 | 7.48 | 0.159 | 13.89 | 0.016 |
| XPO2_HUMAN | CSE1L | Exportin-2 | -2.86 | 0.797 | 7.86 | 0.111 | 9.39 | 0.043 | -1.99 | 0.955 | 11.33 | 0.109 | 13.53 | 0.020 |
| S6A11_HUMAN | SLC6A11 | Sodium- and chloride-dependent GABA transporter 3 | -INF | 0.000 | 8.59 | 0.119 | 10.48 | 0.019 | NA | NA | +INF | 0.000 | +INF | 0.000 |
| SSFA2_HUMAN | SSFA2 | Sperm-specific antigen 2 | -1.34 | 0.919 | 7.11 | 0.222 | 11.98 | 0.018 | 1.11 | 0.982 | 10.60 | 0.158 | 17.86 | 0.024 |
| YKT6_HUMAN | YKT6 | Synaptobrevin homolog YKT6 | -1.75 | 0.921 | 7.04 | 0.310 | 12.03 | 0.044 | -1.91 | 0.973 | 6.46 | 0.076 | 11.04 | 0.005 |
| EHBP1_HUMAN | EHBP1 | EH domain-binding protein 1 | -6.45 | 0.499 | 8.56 | 0.183 | 12.84 | 0.022 | -3.48 | 0.876 | 15.88 | 0.085 | 23.83 | 0.022 |
| LBP_HUMAN | LBP | Lipopolysaccharide-binding protein | 1.24 | 0.919 | 4.39 | 0.278 | 13.17 | 0.009 | -1.12 | 0.980 | 3.15 | 0.367 | 9.47 | 0.020 |
| ROA0_HUMAN | HNRNPA0 | Heterogeneous nuclear ribonucleoprotein A0 | 1.48 | 0.978 | 7.41 | 0.095 | 13.34 | 0.005 | 1.49 | 0.984 | 7.45 | 0.193 | 13.41 | 0.033 |
| SHRM3_HUMAN | SHROOM3 | Protein Shroom3 | -1.98 | 0.906 | 5.03 | 0.251 | 13.61 | 0.022 | 1.44 | 0.968 | 14.30 | 0.039 | 38.67 | 0.000 |
| CCDC6_HUMAN | CCDC6 | Coiled-coil domain-containing protein 6 | 1.09 | 0.979 | 12.15 | 0.149 | 13.73 | 0.044 | 2.31 | 0.978 | 25.71 | 0.036 | 29.05 | 0.010 |
| PARK7_HUMAN | PARK7 | Protein/nucleic acid deglycase DJ-1 | -1.99 | 0.849 | 11.43 | 0.053 | 14.54 | 0.005 | -1.331 | 0.993 | 17.09 | 0.039 | 21.74 | 0.004 |
| TOM40_HUMAN | TOMM40 | Mitochondrial import receptor subunit TOM40 homolog | -12.97 | 0.733 | 2.63 | 0.805 | 15.60 | 0.033 | -4.75 | 0.919 | 7.17 | 0.276 | 42.56 | 0.006 |
| PPIA_HUMAN | PPIA | Peptidyl-prolyl cis-trans isomerase A | -1.81 | 0.816 | 8.13 | 0.078 | 16.58 | 0.003 | -2.14 | 0.955 | 6.87 | 0.154 | 14.01 | 0.020 |
| CTL2_HUMAN | SLC44A2 | Choline transporter-like protein 2 | -4.49 | 0.999 | 8.10 | 0.309 | 16.75 | 0.025 | -3.54 | 0.971 | 10.24 | 0.313 | 21.20 | 0.039 |
| DNJC3_HUMAN | DNAJC3 | DnaJ homolog subfamily C member 3 OS=Homo sapiens | 7.81 | 0.545 | 10.46 | 0.201 | 17.81 | 0.024 | 9.70 | 0.693 | 13.00 | 0.174 | 22.12 | 0.032 |
| SIAE_HUMAN | SIAE | Sialate O-acetylesterase | 2.06 | 0.951 | 11.40 | 0.067 | 19.19 | 0.006 | 2.01 | 0.990 | 11.11 | 0.086 | 18.70 | 0.016 |
| GCC2_HUMAN | GCC2 | GRIP and coiled-coil domain-containing protein 2 | -1.77 | 0.999 | 5.34 | 0.363 | 19.92 | 0.019 | -1.98 | 0.971 | 4.77 | 0.461 | 17.78 | 0.040 |
| DCTN2_HUMAN | DCTN2 | Dynactin subunit 2 | 1.60 | 0.978 | 9.65 | 0.231 | 20.32 | 0.024 | 1.46 | 0.984 | 8.79 | 0.313 | 18.51 | 0.049 |
| RECQ1_HUMAN | RECQL | ATP-dependent DNA helicase Q1 | -1.41 | 0.919 | 3.27 | 0.476 | 21.74 | 0.004 | -1.07 | 0.990 | 4.30 | 0.335 | 28.61 | 0.002 |
| RALY_HUMAN | RALY | RNA-binding protein Raly | 1.11 | 0.997 | 11.57 | 0.093 | 22.98 | 0.002 | -1.09 | 0.992 | 9.55 | 0.134 | 18.98 | 0.008 |
| LRC57_HUMAN | LRRC57 | Leucine-rich repeat-containing protein 57 | 1.64 | 0.999 | 11.04 | 0.014 | 28.60 | 0.000 | -2.08 | 0.971 | 3.24 | 0.332 | 8.40 | 0.010 |
| NCF2_HUMAN | NCF2 | Neutrophil cytosol factor 2 | 1.42 | 0.919 | 4.46 | 0.482 | 29.35 | 0.024 | 1.63 | 0.966 | 5.09 | 0.484 | 33.51 | 0.024 |
| BROX_HUMAN | BROX | BRO1 domain-containing protein BROX | 1.14 | 0.995 | 17.93 | 0.217 | 29.96 | 0.042 | 2.20 | 0.964 | 34.64 | 0.122 | 57.90 | 0.046 |
| PRAF3_HUMAN | ARL6IP5 | PRA1 family protein 3 | 3.59 | 0.942 | 12.63 | 0.306 | 32.81 | 0.033 | 6.44 | 0.919 | 22.66 | 0.211 | 58.88 | 0.028 |
| NIPS1_HUMAN | NIPSNAP1 | Protein NipSnap homolog 1 | -1.68 | 0.965 | 10.78 | 0.115 | 38.57 | 0.002 | -2.11 | 0.938 | 8.60 | 0.260 | 30.79 | 0.010 |
| VPS4A_HUMAN | VPS4A | Vacuolar protein sorting-associated protein 4A | 1.34 | 0.993 | 21.66 | 0.011 | 41.30 | 0.000 | -1.40 | 0.992 | 11.55 | 0.148 | 22.02 | 0.018 |
| AL1L1_HUMAN | ALDH1L1 | Cytosolic 10-formyltetrahydrofolate dehydrogenase | 1.23 | 0.942 | 18.23 | 0.100 | 42.62 | 0.001 | -1.16 | 0.990 | 12.80 | 0.191 | 29.94 | 0.011 |
| STML2_HUMAN | STOML2 | Stomatin-like protein 2, mitochondrial | 1.31 | 0.997 | 19.44 | 0.121 | 44.44 | 0.006 | -1.47 | 0.992 | 10.12 | 0.234 | 23.14 | 0.037 |
| PGBM_HUMAN | HSPG2 | Basement membrane-specific heparan sulfate proteoglycan core protein | 24.22 | 0.713 | 28.76 | 0.028 | 45.90 | 0.003 | 12.90 | 0.825 | 15.32 | 0.154 | 24.45 | 0.036 |
| DJC10_HUMAN | DNAJC10 | DnaJ homolog subfamily C member 10 | 2.55 | 0.999 | 13.10 | 0.363 | 50.58 | 0.025 | 2.26 | 0.971 | 11.60 | 0.318 | 44.81 | 0.029 |
| CO3_HUMAN | C3 | Complement C3 | -2.47 | 0.942 | 15.64 | 0.135 | 56.41 | 0.001 | -2.87 | 0.919 | 13.44 | 0.211 | 48.46 | 0.006 |
| KBTBB_HUMAN | KBTBD11 | Kelch repeat and BTB domain-containing protein 11 | -1.08 | 0.964 | 8.51 | 0.347 | 62.14 | 0.001 | 1.79 | 0.947 | 16.41 | 0.191 | 119.79 | 0.001 |
| HPTR_HUMAN | HPR | Haptoglobin-related protein | -3.72 | 0.942 | 397.79 | 0.075 | 437.23 | 0.008 | -3.01 | 0.971 | 491.59 | 0.102 | 540.34 | 0.019 |
| SUOX_HUMAN | SUOX | Sulfite oxidase, mitochondrial | +INF | - | +INF | 0.000 | +INF | 0.000 | -INF | 0.000 | 2.02 | 0.178 | 2.40 | 0.030 |
| NT5D1_HUMAN | NT5DC1 | 5'-nucleotidase domain-containing protein 1 | +INF | 0.000 | +INF | 0.000 | +INF | 0.000 | -1.17 | 0.964 | 5.28 | 0.094 | 5.13 | 0.046 |

### DE proteins in *IDH-mut* AST

| | | | Fold Change Relative to HC | | | | | | Fold Change Relative to MEN | | | | | |
|---|---|---|---|---|---|---|---|---|---|---|---|---|---|---|
| Protein | | | *IDH-mut* OLI | | *IDH-mut* AST | | *IDH-wt* GBM | | *IDH-mut* OLI | | *IDH-mut* AST | | *IDH-wt* GBM | |
| Accession | Gene Name | Name | FC | *p*-val | FC | *p*-val | FC | *p*-val | FC | *p*-val | FC | *p*-val | FC | *p*-val |
| FXR1_HUMAN | FXR1 | Fragile X mental retardation syndrome-related protein 1 | -1.01 | 0.997 | 8.46 | 0.027 | 4.07 | 0.090 | 1.34 | 0.990 | 11.41 | 0.004 | 5.48 | 0.037 |

| DE proteins in *IDH-mut* OLI | | | | | | | | | | | | | | |
|---|---|---|---|---|---|---|---|---|---|---|---|---|---|---|
| | | | **Fold Change Relative to HC** | | | | | | **Fold Change Relative to MEN** | | | | | |
| **Protein** | | | ***IDH-mut* OLI** | | ***IDH-mut* AST** | | ***IDH-wt* GBM** | | ***IDH-mut* OLI** | | ***IDH-mut* AST** | | ***IDH-wt* GBM** | |
| **Accession** | **Gene Name** | **Name** | **FC** | ***p*-val** | **FC** | ***p*-val** | **FC** | ***p*-val** | **FC** | ***p*-val** | **FC** | ***p*-val** | **FC** | ***p*-val** |
| PDCL3_HUMAN | PDCL3 | Phosducin-like protein 3 | 8.71 | 0.018 | 2.09 | 0.135 | 2.11 | 0.075 | +INF | 0.000 | +INF | 0.000 | +INF | 0.000 |
| TM223_HUMAN | TMEM223 | Transmembrane protein 223 | -INF | 0.000 | 19.68 | 0.229 | 18.34 | 0.245 | -INF | 0.000 | 9.21 | 0.078 | 8.58 | 0.035 |
| NDUS8_HUMAN | NDUFS8 | NADH dehydrogenase [ubiquinone] iron-sulfur protein 8, mitochondrial | -INF | 0.000 | 1.29 | 0.733 | 1.51 | 0.536 | -INF | 0.000 | 5.49 | 0.071 | 6.41 | 0.022 |
| ERCC3_HUMAN | ERCC3 | General transcription and DNA repair factor IIH helicase subunit XPB | -INF | 0.000 | 10.34 | 0.315 | 13.89 | 0.077 | -INF | 0.000 | 17.33 | 0.158 | 23.28 | 0.048 |
| SNTA1_HUMAN | SNTA1 | Alpha-1-syntrophin | -INF | 0.000 | -1.64 | 0.795 | -1.75 | 0.627 | -INF | 0.000 | 6.63 | 0.098 | 6.20 | 0.040 |
| FA50A_HUMAN | FAM50A | Protein FAM50A | -INF | 0.000 | 1.84 | 0.597 | 1.68 | 0.479 | -INF | 0.000 | 12.60 | 0.064 | 11.46 | 0.032 |
| TRIPB_HUMAN | TRIP11 | Thyroid receptor-interacting protein 11 | -INF | 0.000 | 2.15 | 0.485 | 2.52 | 0.163 | -INF | 0.000 | 3.61 | 0.101 | 4.23 | 0.024 |
| SV2A_HUMAN | SV2A | Synaptic vesicle glycoprotein 2A | -INF | 0.000 | 10.84 | 0.110 | 10.70 | 0.036 | -INF | 0.000 | 7.19 | 0.352 | 7.10 | 0.237 |
| TM209_HUMAN | TMEM209 | Transmembrane protein 209 | -INF | 0.000 | -12.49 | 0.115 | -23.00 | 0.012 | -INF | 0.000 | -2.99 | 0.559 | -5.29 | 0.174 |
| LNP_HUMAN | LNPK | Endoplasmic reticulum junction formation protein lunapark | -INF | 0.000 | 6.03 | 0.133 | 9.26 | 0.014 | -INF | 0.000 | 5.16 | 0.304 | 7.91 | 0.126 |
| ARMX1_HUMAN | ARMCX1 | Armadillo repeat-containing X-linked protein 1 | -INF | 0.000 | 2.47 | 0.277 | 1.99 | 0.226 | -INF | 0.000 | 4.80 | 0.102 | 3.87 | 0.044 |

| DE proteins in *IDH-wt* GBM and *IDH-mut* AST | | | | | | | | | | | | | | |
|---|---|---|---|---|---|---|---|---|---|---|---|---|---|---|
| | | | **Fold Change Relative to HC** | | | | | | **Fold Change Relative to MEN** | | | | | |
| **Protein** | | | ***IDH*-mut OLI** | | ***IDH*-mut AST** | | ***IDH*-wt GBM** | | ***IDH*-mut OLI** | | ***IDH*-mut AST** | | ***IDH*-wt GBM** | |
| **Accession** | **Gene Name** | **Name** | **FC** | ***p*-val** | **FC** | ***p*-val** | **FC** | ***p*-val** | **FC** | ***p*-val** | **FC** | ***p*-val** | **FC** | ***p*-val** |
| IDH3A_HUMAN | IDH3A | Isocitrate dehydrogenase [NAD] subunit alpha, mitochondrial | -1.18 | 0.937 | 5.90 | 0.027 | 6.89 | 0.004 | 1.79 | 0.977 | 12.46 | 0.001 | 14.54 | 0.000 |
| GPC1_HUMAN | GPC1 | Glypican-1 | 2.41 | 0.891 | 14.92 | 0.028 | 37.70 | 0.000 | 2.98 | 0.977 | 18.46 | 0.016 | 46.66 | 0.000 |
| TBCD_HUMAN | TBCD | Tubulin-specific chaperone D | 1.47 | 0.985 | 26.94 | 0.006 | 22.77 | 0.002 | 1.61 | 0.981 | 29.54 | 0.040 | 24.97 | 0.016 |
| BBS2_HUMAN | BBS2 | Bardet-Biedl syndrome 2 protein | -1.30 | 0.994 | 20.21 | 0.006 | 16.09 | 0.003 | 2.38 | 0.938 | 62.21 | 0.002 | 49.54 | 0.002 |
| KAD2_HUMAN | AK2 | Adenylate kinase 2, mitochondrial | -INF | 0.000 | 3106.54 | 0.001 | 613.57 | 0.003 | NA | NA | +INF | 0.000 | +INF | 0.000 |
| MMP25_HUMAN | MMP25 | Matrix metalloproteinase-25 | -327.67 | 0.057 | -116.91 | 0.025 | -127.54 | 0.013 | +INF | 0.000 | +INF | 0.000 | +INF | 0.000 |
| PRDX5_HUMAN | PRDX5 | Peroxiredoxin-5, mitochondrial | 1.36 | 0.900 | 9.09 | 0.039 | 8.56 | 0.009 | +INF | 0.000 | +INF | 0.000 | +INF | 0.000 |
| HEM3_HUMAN | HMBS | Porphobilinogen deaminase | -5.46 | 0.887 | -280.61 | 0.037 | -243.35 | 0.005 | -13.92 | 0.977 | -715.36 | 0.015 | -620.47 | 0.001 |
| RL22_HUMAN | RPL22 | 60S ribosomal protein L22 | -7.50 | 0.729 | -20.07 | 0.027 | -20.56 | 0.005 | -29.58 | 0.976 | -79.12 | 0.001 | -81.06 | 0.000 |

| DE proteins in *IDH-wt* GBM and *IDH-mut* OLI | | | | | | | | | |
|---|---|---|---|---|---|---|---|---|---|
| | | | **Fold Change Relative to HC** | | | | | | **Fold Change Relative to MEN** |

| Protein | | | *IDH-mut* OLI | | *IDH-mut* AST | | *IDH-wt* GBM | | *IDH-mut* OLI | | *IDH-mut* AST | | *IDH-wt* GBM | |
|---|---|---|---|---|---|---|---|---|---|---|---|---|---|---|
| Accession | Gene Name | Name | FC | *p*-val | FC | *p*-val | FC | *p*-val | FC | *p*-val | FC | *p*-val | FC | *p*-val |
| TIPIN_HUMAN | TIPIN | TIMELESS-interacting protein | -INF | 0 | 2.23 | 0.580 | -22.34 | 0.012 | -INF | 0.000 | 1.70 | 0.865 | -29.34 | 0.004 |
| **DE proteins in Glioma (*IDH-wt* GBM, *IDH-mut* AST and *IDH-mut* OLI)** | | | | | | | | | | | | | | |
| | | | **Fold Change Relative to HC** | | | | | | **Fold Change Relative to MEN** | | | | | |
| **Protein** | | | ***IDH-mut* OLI** | | ***IDH-mut* AST** | | ***IDH-wt* GBM** | | ***IDH-mut* OLI** | | ***IDH-mut* AST** | | ***IDH-wt* GBM** | |
| **Accession** | **Gene Name** | **Name** | **FC** | ***p*-val** | **FC** | ***p*-val** | **FC** | ***p*-val** | **FC** | ***p*-val** | **FC** | ***p*-val** | **FC** | ***p*-val** |
| EBP2_HUMAN | EBNA1BP2 | Probable rRNA-processing protein EBP2 | +INF | 0 | +INF | 0 | +INF | 0 | +INF | 0 | +INF | 0 | +INF | 0 |
| NIBAN_HUMAN | FAM129A | Protein Niban | +INF | 0 | +INF | 0 | +INF | 0 | +INF | 0 | +INF | 0 | +INF | 0 |

**Table 5 – Glioma-specific protein changes in plasma EVs associated with glioma grade.** DE proteins relative to both healthy and non-glioma controls. Proteins that are restricted to glioma samples are denoted by '+INF' and proteins that are restricted to controls (HC or MEN) are denoted by '-INF'.

| | | | DE Proteins in Grade IV Glioma Plasma EVs | | | | | | | | | | | |
|---|---|---|---|---|---|---|---|---|---|---|---|---|---|---|
| | | | Fold Change Relative to HC | | | | | | Fold Change Relative to MEN | | | | | |
| Protein | | | Grade II | | Grade III | | Grade IV | | Grade II | | Grade III | | Grade IV | |
| Accession | Gene name | Name | FC | *p*-val | FC | *p*-val | FC | *p*-val | FC | *p*-val | FC | *p*-val | FC | *p*-val |
| 1A03_HUMAN | HLA-A | HLA class I histocompatibility antigen, A-3 alpha chain | -209.36 | 0.400 | -23.61 | 0.911 | -539.20 | 0.007 | -238.67 | 0.463 | -26.91 | 0.860 | -614.68 | 0.009 |
| SAP_HUMAN | PSAP | Prosaposin | -92.55 | 0.400 | -134.12 | 0.273 | -276.91 | 0.007 | -41.62 | 0.610 | -60.32 | 0.517 | -124.53 | 0.022 |
| HEM3_HUMAN | HMBS | Porphobilinogen deaminase | -103.83 | 0.400 | -40.73 | 0.901 | -257.61 | 0.007 | -264.71 | 0.283 | -103.82 | 0.517 | -656.74 | 0.003 |
| HEM4_HUMAN | UROS | Uroporphyrinogen-III synthase | -42.94 | 0.730 | -5.20 | 0.911 | -232.81 | 0.007 | -273.09 | 0.283 | -33.04 | 0.658 | -1480.5 | 0.001 |
| CO6_HUMAN | C6 | Complement component C6 | -1.06 | 0.991 | -8.97 | 0.911 | -223.87 | 0.009 | 1.73 | 0.980 | -4.87 | 0.996 | -121.65 | 0.038 |
| DQB1_HUMAN | HLA-DQB1 | HLA class II histocompatibility antigen, DQ beta 1 chain | -6.93 | 0.879 | -18.11 | 0.901 | -117.25 | 0.007 | -18.15 | 0.681 | -47.42 | 0.517 | -307.01 | 0.001 |
| ACYP2_HUMAN | ACYP2 | Acylphosphatase-2 | -8.81 | 0.902 | -7.94 | 0.911 | -90.66 | 0.026 | -25.51 | 0.733 | -23.00 | 0.849 | -262.66 | 0.007 |
| ENOG_HUMAN | ENO2 | Gamma-enolase | -9.78 | 0.778 | -2.39 | 0.976 | -42.95 | 0.044 | -15.43 | 0.733 | -3.77 | 0.996 | -67.73 | 0.027 |
| NUF2_HUMAN | NUF2 | Kinetochore protein Nuf2 | -2.40 | 0.792 | -3.28 | 0.487 | -24.62 | 0.001 | -1.65 | 0.904 | -2.25 | 0.793 | -16.87 | 0.009 |
| EPS8_HUMAN | EPS8 | Epidermal growth factor receptor kinase substrate 8 | -15.12 | 0.304 | -6.55 | 0.713 | -23.05 | 0.013 | -18.01 | 0.280 | -7.80 | 0.642 | -27.45 | 0.030 |
| TIPIN_HUMAN | TIPIN | TIMELESS-interacting protein | 1.61 | 0.964 | 2.62 | 0.759 | -22.34 | 0.018 | 1.23 | 0.970 | 1.99 | 0.867 | -29.34 | 0.008 |
| RL22_HUMAN | RPL22 | 60S ribosomal protein L22 | -10.37 | 0.527 | -28.55 | 0.075 | -20.77 | 0.005 | -40.87 | 0.099 | -112.55 | 0.010 | -81.88 | 0.000 |
| STIM1_HUMAN | STIM1 | Stromal interaction molecule 1 | -3.10 | 0.701 | -6.37 | 0.713 | -13.92 | 0.013 | -2.25 | 0.801 | -4.62 | 0.706 | -10.09 | 0.049 |
| ITA7_HUMAN | ITGA7 | Integrin alpha-7 | -7.46 | 0.519 | -2.46 | 0.831 | -11.81 | 0.024 | -8.82 | 0.526 | -2.91 | 0.876 | -13.97 | 0.037 |
| ACSF2_HUMAN | ACSF2 | Acyl-CoA synthetase family member 2, mitochondrial | -2.38 | 0.784 | -4.54 | 0.439 | -11.70 | 0.005 | -3.73 | 0.518 | -7.13 | 0.204 | -18.37 | 0.002 |
| SYTM_HUMAN | TARS2 | Threonine--tRNA ligase, mitochondrial | -3.89 | 0.652 | -3.23 | 0.672 | -7.19 | 0.018 | -4.05 | 0.936 | -3.36 | 0.787 | -7.47 | 0.042 |
| SVIP_HUMAN | SVIP | Small VCP/p97-interacting protein | -1.24 | 0.979 | -1.64 | 0.841 | -7.10 | 0.031 | -1.41 | 0.971 | -1.88 | 0.861 | -8.10 | 0.026 |
| BD1L1_HUMAN | BOD1L1 | Biorientation of chromosomes in cell division protein 1-like 1 | -9.16 | 0.549 | 1.29 | 0.949 | -6.94 | 0.031 | -14.59 | 0.180 | -1.24 | 0.997 | -11.05 | 0.014 |
| HSP7E_HUMAN | HSPA14 | Heat shock 70 kDa protein 14 | -2.11 | 0.647 | -2.75 | 0.672 | -5.69 | 0.003 | -2.20 | 0.615 | -2.87 | 0.427 | -5.94 | 0.005 |
| PCNT_HUMAN | PCNT | Pericentrin | -5.50 | 0.332 | -2.47 | 0.827 | -5.67 | 0.041 | -11.70 | 0.121 | -5.25 | 0.534 | -12.07 | 0.009 |
| NGLY1_HUMAN | NGLY1 | Peptide-N(4)-(N-acetyl-beta-glucosaminyl)asparagine amidase | -3.52 | 0.779 | -5.71 | 0.272 | -5.64 | 0.033 | -5.49 | 0.231 | -8.91 | 0.101 | -8.81 | 0.008 |
| WDR61_HUMAN | WDR61 | WD repeat-containing protein 61 | -3.52 | 0.253 | -2.16 | 0.485 | -4.35 | 0.004 | -3.81 | 0.263 | -2.34 | 0.461 | -4.71 | 0.007 |
| TRI14_HUMAN | TRIM14 | Tripartite motif-containing protein 14 | -1.56 | 0.869 | -2.25 | 0.713 | -4.20 | 0.019 | -1.87 | 0.801 | -2.68 | 0.650 | -5.01 | 0.031 |
| K154L_HUMAN | KIAA1549L | UPF0606 protein KIAA1549L | -2.29 | 0.821 | -2.05 | 0.733 | -3.38 | 0.034 | -8.98 | 0.006 | -8.06 | 0.004 | -13.28 | 0.000 |
| DGKA_HUMAN | DGKA | Diacylglycerol kinase alpha | -1.23 | 0.925 | -2.19 | 0.716 | -3.35 | 0.028 | -1.37 | 0.888 | -2.45 | 0.714 | -3.73 | 0.032 |
| SUOX_HUMAN | SUOX | Sulfite oxidase, mitochondrial | INF | 0 | NA | NA | INF | 0 | 2.05 | 0.400 | -INF | 0 | 2.33 | 0.042 |
| TOPK_HUMAN | PBK | Lymphokine-activated killer T-cell-originated protein kinase | 0.02 | 0.779 | 0.0 1 | 0.289 | 0.01 | 0.033 | INF | 0 | INF | 0 | INF | 0 |
| TLK2_HUMAN | TLK2 | Serine/threonine-protein kinase tousled-like 2 | -INF | 0 | 2.06 | 0.776 | 0.18 | 0.018 | NA | NA | INF | 0 | INF | 0 |
| PRDX5_HUMAN | PRDX5 | Peroxiredoxin-5, mitochondrial | 4.52 | 0.631 | 5.15 | 0.7166 | 8.62 | 0.026 | INF | 0 | INF | 0 | INF | 0 |
| S6A11_HUMAN | SLC6A11 | Sodium- and chloride-dependent GABA transporter 3 | 10.29 | 0.387 | 5.82 | 0.663 | 10.49 | 0.026 | INF | 0 | INF | 0 | INF | 0 |
| API5_HUMAN | API5 | Apoptosis inhibitor 5 | 1.03 | 0.990 | 3.08 | 0.487 | 5.51 | 0.042 | 1.26 | 0.941 | 3.78 | 0.461 | 6.75 | 0.010 |
| APC1_HUMAN | ANAPC1 | Anaphase-promoting complex subunit 1 | 7.34 | 0.253 | 3.39 | 0.454 | 5.89 | 0.007 | 9.01 | 0.312 | 4.17 | 0.461 | 7.24 | 0.021 |
| TCPH_HUMAN | CCT7 | T-complex protein 1 subunit eta | 1.949 | 0.818 | 2.368 | 0.759 | 7.204 | 0.011 | 1.79 | 0.970 | 2.17 | 0.867 | 6.61 | 0.042 |
| WIPI3_HUMAN | WDR45B | WD repeat domain phosphoinositide-interacting protein 3 | 5.028 | 0.384 | 2.023 | 0.917 | 7.383 | 0.017 | 7.368 | 0.231 | 2.97 | 0.752 | 10.82 | 0.016 |
| IDH3A_HUMAN | IDH3A | Isocitrate dehydrogenase [NAD] subunit alpha, | 3.40 | 0.596 | 2.55 | 0.675 | 7.70 | 0.005 | 7.17 | 0.126 | 5.39 | 0.192 | 16.25 | 0.000 |

| | | | | | | | | | | | | | | |
|---|---|---|---|---|---|---|---|---|---|---|---|---|---|---|
| | | mitochondrial | | | | | | | | | | | | |
| CPTP_HUMAN | CPTP | Ceramide-1-phosphate transfer protein | 2.98 | 0.501 | 5.20 | 0.162 | 7.87 | 0.001 | 2.17 | 0.584 | 3.78 | 0.508 | 5.71 | 0.012 |
| ANLN_HUMAN | ANLN | Anillin | 4.52 | 0.904 | 1.62 | 0.938 | 10.49 | 0.016 | 6.69 | 0.664 | 2.39 | 0.808 | 15.51 | 0.008 |
| STAU1_HUMAN | STAU1 | Double-stranded RNA-binding protein Staufen homolog 1 | 3.95 | 0.580 | 1.90 | 0.976 | 10.54 | 0.041 | 5.99 | 0.449 | 2.88 | 0.854 | 16.01 | 0.023 |
| SSFA2_HUMAN | SSFA2 | Sperm-specific antigen 2 | 7.92 | 0.631 | 3.07 | 0.777 | 11.87 | 0.023 | 11.81 | 0.463 | 4.58 | 0.759 | 17.70 | 0.032 |
| MYCB2_HUMAN | MYCBP2 | E3 ubiquitin-protein ligase MYCBP2 | 1.17 | 0.960 | 2.14 | 0.976 | 12.05 | 0.048 | 1.92 | 0.878 | 3.49 | 0.816 | 19.70 | 0.019 |
| ROA0_HUMAN | HNRNPA0 | Heterogeneous nuclear ribonucleoprotein A0 | 10.10 | 0.304 | 2.61 | 0.713 | 13.05 | 0.003 | 10.15 | 0.379 | 2.62 | 0.876 | 13.11 | 0.037 |
| SHRM3_HUMAN | SHROOM3 | Protein Shroom3 | 1.66 | 0.931 | 3.22 | 0.754 | 13.33 | 0.028 | 4.72 | 0.719 | 9.16 | 0.257 | 37.88 | 0.001 |
| LBP_HUMAN | LBP | Lipopolysaccharide-binding protein | 4.22 | 0.631 | 1.91 | 0.786 | 13.63 | 0.003 | 3.04 | 0.641 | 1.38 | 0.978 | 9.80 | 0.008 |
| PPIA_HUMAN | PPIA | Peptidyl-prolyl cis-trans isomerase A | 10.43 | 0.294 | 3.97 | 0.745 | 14.69 | 0.011 | 8.81 | 0.713 | 3.36 | 0.816 | 12.41 | 0.040 |
| PARK7_HUMAN | PARK7 | Protein/nucleic acid deglycase DJ-1 OS=Homo sapiens OX=9606 GN=PE=1 SV=2 | 7.59 | 0.481 | 3.10 | 0.759 | 15.26 | 0.009 | 11.35 | 0.391 | 4.64 | 0.787 | 22.83 | 0.008 |
| TOM40_HUMAN | TOMM40 | Mitochondrial import receptor subunit TOM40 homolog | 1.197 | 0.960 | -1.29 | 0.991 | 15.28 | 0.043 | 3.26 | 0.779 | 2.13 | 0.854 | 41.69 | 0.010 |
| BBS2_HUMAN | BBS2 | Bardet-Biedl syndrome 2 protein | 10.55 | 0.304 | 9.97 | 0.430 | 17.52 | 0.007 | 32.48 | 0.167 | 30.70 | 0.225 | 53.92 | 0.008 |
| DNJC3_HUMAN | DNAJC3 | DnaJ homolog subfamily C member 3 | 11.81 | 0.519 | 7.30 | 0.713 | 18.13 | 0.017 | 14.67 | 0.427 | 9.07 | 0.618 | 22.53 | 0.031 |
| CTL2_HUMAN | SLC44A2 | Choline transporter-like protein 2 | 3.10 | 0.968 | 1.60 | 0.928 | 19.31 | 0.026 | 3.92 | 0.939 | 2.03 | 0.895 | 24.44 | 0.038 |
| SIAE_HUMAN | SIAE | Sialate O-acetylesterase | 7.09 | 0.391 | 8.03 | 0.419 | 19.88 | 0.006 | 6.91 | 0.538 | 7.82 | 0.461 | 19.37 | 0.015 |
| DCTN2_HUMAN | DCTN2 | Dynactin subunit 2 | 8.71 | 0.519 | 3.99 | 0.713 | 20.24 | 0.019 | 7.93 | 0.587 | 3.63 | 0.876 | 18.44 | 0.049 |
| RECQ1_HUMAN | RECQL | ATP-dependent DNA helicase Q1 | 1.88 | 0.904 | 1.90 | 0.885 | 20.25 | 0.005 | 2.47 | 0.784 | 2.50 | 0.937 | 26.65 | 0.003 |
| IVD_HUMAN | IVD | Isovaleryl-CoA dehydrogenase, mitochondrial | 2.53 | 0.870 | 17.09 | 0.466 | 20.40 | 0.026 | 1.58 | 0.893 | 10.65 | 0.439 | 12.71 | 0.038 |
| GCC2_HUMAN | GCC2 | GRIP and coiled-coil domain-containing protein 2 | 3.36 | 0.914 | 1.70 | 0.928 | 20.72 | 0.011 | 3.00 | 0.939 | 1.52 | 0.919 | 18.50 | 0.038 |
| BTF3_HUMAN | BTF3 | Transcription factor BTF3 | 34.61 | 0.183 | 6.56 | 0.716 | 21.20 | 0.006 | 20.08 | 0.427 | 3.80 | 0.832 | 12.30 | 0.032 |
| MCM3_HUMAN | MCM3 | DNA replication licensing factor MCM3 | 5.50 | 0.631 | 4.24 | 0.730 | 21.36 | 0.006 | 2.33 | 0.888 | 1.80 | 0.956 | 9.07 | 0.049 |
| EIF3I_HUMAN | EIF3I | Eukaryotic translation initiation factor 3 subunit I | 2.03 | 0.881 | 5.36 | 0.713 | 22.33 | 0.017 | 2.19 | 0.848 | 5.77 | 0.791 | 24.03 | 0.031 |
| TBCD_HUMAN | TBCD | Tubulin-specific chaperone D | 13.48 | 0.304 | 10.18 | 0.430 | 22.99 | 0.004 | 14.78 | 0.721 | 11.16 | 0.631 | 25.21 | 0.042 |
| RALY_HUMAN | RALY | RNA-binding protein Raly | 7.12 | 0.632 | 5.04 | 0.616 | 25.17 | 0.001 | 5.88 | 0.528 | 4.16 | 0.660 | 20.78 | 0.010 |
| PAIRB_HUMAN | SERBP1 | Plasminogen activator inhibitor 1 RNA-binding protein | 2.15 | 0.968 | 5.44 | 0.794 | 25.71 | 0.032 | 2.95 | 0.939 | 7.47 | 0.669 | 35.29 | 0.039 |
| LRC57_HUMAN | LRRC57 | Leucine-rich repeat-containing protein 57 | 2.75 | 0.874 | 11.71 | 0.020 | 29.59 | 0.000 | -1.24 | 0.958 | 3.44 | 0.641 | 8.69 | 0.007 |
| NCF2_HUMAN | NCF2 | Neutrophil cytosol factor 2 | 7.29 | 0.752 | 1.40 | 0.936 | 31.40 | 0.018 | 8.32 | 0.573 | 1.60 | 0.978 | 35.85 | 0.015 |
| GPC1_HUMAN | GPC1 | Glypican-1 | 12.95 | 0.387 | 7.15 | 0.584 | 38.24 | 0.000 | 16.03 | 0.183 | 8.84 | 0.335 | 47.33 | 0.000 |
| PRAF3_HUMAN | ARL6IP5 | PRA1 family protein 3 | 3.54 | 0.727 | 10.49 | 0.602 | 39.90 | 0.010 | 6.36 | 0.558 | 18.82 | 0.534 | 71.60 | 0.010 |
| NIPS1_HUMAN | NIPSNAP1 | Protein NipSnap homolog 1 | 4.25 | 0.714 | 4.77 | 0.672 | 39.97 | 0.003 | 3.39 | 0.970 | 3.81 | 0.840 | 31.91 | 0.011 |
| VPS4A_HUMAN | VPS4A | Vacuolar protein sorting-associated protein 4A | 7.38 | 0.632 | 19.11 | 0.161 | 41.89 | 0.001 | 3.94 | 0.634 | 10.19 | 0.460 | 22.34 | 0.029 |
| AL1L1_HUMAN | ALDH1L1 | Cytosolic 10-formyltetrahydrofolate dehydrogenase | 10.65 | 0.332 | 10.22 | 0.602 | 42.95 | 0.002 | 7.48 | 0.449 | 7.18 | 0.752 | 30.17 | 0.012 |
| DJC10_HUMAN | DNAJC10 | DnaJ homolog subfamily C member 10 | 9.22 | 0.783 | 6.65 | 0.794 | 49.63 | 0.026 | 8.17 | 0.863 | 5.89 | 0.671 | 43.97 | 0.031 |
| PGBM_HUMAN | HSPG2 | Basement membrane-specific heparan sulfate proteoglycan core protein | 16.12 | 0.294 | 23.06 | 0.227 | 52.73 | 0.002 | 8.59 | 0.718 | 12.29 | 0.552 | 28.09 | 0.035 |
| CO3_HUMAN | C3 | Complement C3 | 6.14 | 0.503 | 3.81 | 0.887 | 59.06 | 0.002 | 5.28 | 0.584 | 3.27 | 0.854 | 50.74 | 0.010 |
| KBTBB_HUMAN | KBTBD11 | Kelch repeat and BTB domain-containing protein 11 | 1.28 | 0.942 | 5.97 | 0.769 | 66.95 | 0.001 | 2.46 | 0.841 | 11.50 | 0.534 | 129.05 | 0.000 |
| HPTR_HUMAN | HPR | Haptoglobin-related protein | 152.91 | 0.332 | 81.13 | 0.602 | 502.29 | 0.010 | 188.98 | 0.353 | 100.26 | 0.573 | 620.73 | 0.022 |

**DE proteins in Grade II Glioma Plasma EVs**

| | | | Fold Change Relative to HC | | | | | | Fold Change Relative to MEN | | | | | |
|---|---|---|---|---|---|---|---|---|---|---|---|---|---|---|
| Protein | | | Grade II | | Grade III | | Grade IV | | Grade II | | Grade III | | Grade IV | |
| **Accession** | **Gene name** | **Name** | **FC** | ***p*-val** | **FC** | ***p*-val** | **FC** | ***p*-val** | **FC** | ***p*-val** | **FC** | ***p*-val** | **FC** | ***p*-val** |
| STX10_HUMAN | STX10 | Syntaxin-10 | INF | 0 | INF | 0 | INF | 0 | 55.74 | 0.003 | 6.39 | 0.2504 | 3.50 | 0.191 |
| METL5_HUMAN | METTL5 | Methyltransferase-like protein 5 | -INF | 0 | -2.74 | 0.885 | 5.19 | 0.230 | -INF | 0 | -11.45 | 0.394 | 1.24 | 0.885 |

### DE proteins in Grade II and Grade IV Glioma Plasma EVs

| | | | Fold Change Relative to HC | | | | | | Fold Change Relative to MEN | | | | | |
|---|---|---|---|---|---|---|---|---|---|---|---|---|---|---|
| Protein | | | Grade II | | Grade III | | Grade IV | | Grade II | | Grade III | | Grade IV | |
| **Accession** | **Gene name** | **Name** | **FC** | ***p*-val** | **FC** | ***p*-val** | **FC** | ***p*-val** | **FC** | ***p*-val** | **FC** | ***p*-val** | **FC** | ***p*-val** |
| LRRC7_HUMAN | LRRC7 | Leucine-rich repeat-containing protein 7 | 7.46 | 0.020 | 1.37 | 0.881 | 3.03 | 0.007 | Inf | 0 | Inf | 0 | Inf | 0 |
| PELO_HUMAN | PELO | Protein pelota homolog | -14.5 | 0.005 | -2.48 | 0.672 | -7.59 | 0.003 | -12.62 | 0.021 | -2.16 | 0.840 | -6.60 | 0.008 |
| MMP25_HUMAN | MMP25 | Matrix metalloproteinase-25 | -189.7 | 0.013 | -96.23 | 0.094 | -124.52 | 0.0138 | Inf | 0 | Inf | 0 | Inf | 0 |

### DE proteins in Grade II, Grade III and Grade IV Glioma Plasma EVs

| | | | Fold Change Relative to HC | | | | | | Fold Change Relative to MEN | | | | | |
|---|---|---|---|---|---|---|---|---|---|---|---|---|---|---|
| Protein | | | Grade II | | Grade III | | Grade IV | | Grade II | | Grade III | | Grade IV | |
| **Accession** | **Gene name** | **Name** | **FC** | ***p*-val** | **FC** | ***p*-val** | **FC** | ***p*-val** | **FC** | ***p*-val** | **FC** | ***p*-val** | **FC** | ***p*-val** |
| CYB5_HUMAN | CYB5A | Cytochrome b5 | INF | 0 | INF | 0 | INF | 0 | INF | 0 | INF | 0 | INF | 0 |
| KAD2_HUMAN | AK2 | Adenylate kinase 2, mitochondrial | 3791.6 | 0.0372 | 2986.2 | 0.0107 | 705.85 | 0.0106 | INF | 0 | INF | 0 | INF | 0 |
| GOT1B_HUMAN | GOLT1B | Vesicle transport protein GOT1B | INF | 0 | INF | 0 | INF | 0 | INF | 0 | INF | 0 | INF | 0 |
